# LRRK2 G2019S mutation incites increased cell-intrinsic neutrophil effector functions and intestinal inflammation in a model of infectious colitis

**DOI:** 10.1101/2024.11.26.625468

**Authors:** Jessica Pei, Nathalia L. Oliveira, Sherilyn J. Recinto, Alexandra Kazanova, Celso M. Queiroz-Junior, Ziyi Li, Katalina Couto, Susan Westfall, Irah L. King, Camila T. Ribeiro, Austen J. Milnerwood, Michel Desjardins, Ajitha Thanabalasuriar, Jo Ann Stratton, Samantha Gruenheid

**Author notes:** Equal contribution (Jessica Pei, Nathalia L. Oliveira).

## Abstract

Parkinson’s Disease (PD) is a progressive, neurodegenerative disorder characterised by motor and non-motor symptoms. Emerging evidence suggests a link between PD and gastrointestinal dysfunction. Constipation is frequently observed years prior to development of motor dysfunction in PD, and people with inflammatory bowel disease (IBD) are more likely to develop PD. Mutations in the leucine-rich repeat kinase 2 gene (*LRRK2*) account for approximately 1% of all PD cases and are associated with increased risk for IBD. Among them, LRRK2 Gly2019Ser (G2019S), located within the kinase domain, is the most common PD-associated mutation and increases kinase activity. It is unknown how LRRK2 mutation affects susceptibility to intestinal inflammation or pathogenesis of PD. Using single cell RNA sequencing (scRNAseq), we demonstrate that LRRK2 G2019S mutation promotes a dysregulated gene profile, especially within neutrophil, monocyte and γδ T cell populations, following *Citrobacter rodentium* infection in mice. Transcriptionally, LRRK2 G2019S neutrophils have a greater pro-inflammatory type I and II IFN response compared to those of WT mice. This is accompanied by an increase in neutrophil numbers in the lamina propria in LRRK2 G2019S mice. We also uncover cell-intrinsic functional defects in LRRK2 G2019S neutrophils, including increased chemotaxis, degranulation and neutrophil extracellular traps (NETosis) formation. Increased neutrophil infiltration is associated with an upregulation in Th17 immune responses, which may together contribute to the observed increase in colon pathology during infection. These findings increase our understanding of the role of PD-associated genes in immune cells and their contribution to immune dysregulation. Understanding the early perturbations driven by the LRRK2 G2019S mutation in gastrointestinal pathology may facilitate the development of biomarkers for early diagnosis and intervention in PD.

## Introduction

Parkinson’s disease (PD) is a neurological disease associated with aging. It is the second most prevalent neurodegenerative disorder, after Alzheimer’s disease (Aarsland et al., 2021), and is predicted to affect 14 million individuals by 2040 (Dorsey et al., 2018). Hallmark symptoms of PD include bradykinesia, rigidity, and resting tremor which are linked to the loss of dopaminergic neurons in the substantia nigra pars compacta region of the brain (Aarsland et al., 2021). The average age of PD diagnosis is about 60 years of age, when it is estimated that up to 70% of dopaminergic neurons have already been destroyed (Dauer & Przedborski, 2003). Currently accessible treatments can alleviate some PD motor symptoms, but these therapies loose effectiveness over time and do not target the wide range of other symptoms. (Dauer & Przedborski, 2003). PD is associated with a diverse range of non-motor symptoms that significantly increase the overall burden of the condition. This includes sleep disturbances, hyposmia, and gastrointestinal issues such as constipation, dysphagia, and small intestinal bacterial overgrowth, which can manifest decades before the disease progresses to motor dysfunction (Fang et al., 2024; Tansey et al., 2022). There is therefore considerable interest in the field to increase our understanding of the early, pre-motor pathophysiology of PD, essential for better early diagnosis and management of the disease.

Of note, several studies have indicated that PD is more common in people with IBD and those with previous evidence of intestinal infection (Nerius et al., 2020; Zhu et al., 2022), suggesting that intestinal inflammation can contribute to PD pathogenesis. Mutations in the leucine-rich repeat kinase 2 gene (LRRK2) are responsible for approximately 1% of all PD cases (Volta et al., 2017) and increased risk for IBD (Kuhlmann & Milnerwood, 2020). In contrast to the brain, LRRK2 is extensively expressed in peripheral organs such as the lungs, spleen, and kidneys (Wang et al., 2023). LRRK2 is highly expressed in a variety of immune cells, including, neutrophils, macrophages, monocytes, and B cells (Wang et al., 2023), suggesting a possible immune mediated effect of LRRK2 mutation. Indeed, studies have shown that LRRK2 may play a role in controlling inflammation and pathogen defense in bacterial diseases. LRRK2 has been implicated in increased susceptibility to *Listeria monocytogenes* (Zhang et al., 2015) and in the control of *Salmonella typhimurium* (Liu et al., 2017; Shutinoski et al., 2019). Fang et al. recently demonstrated that male mice carrying the LRRK2 G2019S mutation had increased severity of intestinal inflammation following dextran sodium sulfate (DSS)-induced colitis with changes in the development and severity of PD-related neuropathological and behavioral signs (Fang et al., 2024). Others have found the relationship between LRRK2 mutations and the cGAS-STING pathway, including the systemic production of type I IFNs (Weindel et al., 2020).

Here, we sought to systematically investigate the effects of the LRRK2 G2019S mutation at steady state and in the very early response to intestinal infection. Using the *C. rodentium* model of self-limiting infectious colitis, we show that LRRK2 G2019S promotes increased colon immunopathology following infection, which is associated with an influx of neutrophils and differential gene regulation in immune cells including monocytes, neutrophils and γδT cells. Furthermore, we demonstrate a cell-intrinsic role for LRRK2 G2019S in increasing neutrophil migration, degranulation and NETosis. We also observed an upregulation of Th17 immune responses, which may together contribute to the observed increased colon pathology. Collectively, these findings increase our understanding of the role of PD-associated genes in immune cells and their contribution to immune dysregulation, which could contribute to the development of pharmacological targets and biomarkers for an easier detection and intervention in PD.

## Methods

### Ethics statement and mice

All animal experiments were performed in compliance to the guidelines and conditions specified by the Canadian Council on Animal Care and were approved by the animal care committee of McGill University, animal use protocol number MCGL-5009. Male and female LRRK2 homozygous knock-in G2019S (Yue et al., 2015), and their respective littermate wild-type (WT) mice were used. Mice were bred and maintained under specific pathogen free conditions at the animal facility of McGill University. Sample sizes were based on previous similarly designed experiments from our research group, with five mice per experimental group. Exact mouse numbers for each experiment are included in the figure legends. Mice were randomly assigned to different experiments. G2019S mice were co-housed with WT littermates for all experiments. Animal studies were not blinded. Histopathology scoring, multiplex cytokines and chemokines, and neutrophil migration counting were conducted blindly.

### Citrobacter rodentium infection

The chloramphenicol-resistant *C. rodentium* strain DBS100 (Popov et al., 2019) was used in this study for all *in vivo* and *in vitro* experiments. *C. rodentium* inocula were prepared by culturing bacteria overnight at 37° C, 5% CO2 in Luria-Bertani broth (LB). Bacteria were washed and resuspended in LB and the bacterial density was determined by optical density (O.D.) measured at 600 nm with a spectrophotometer. Mice were inoculated by oral gavage with approximately 10^9^ CFU of *C. rodentium* diluted in 100 μl of LB. Control mice received 100 μl of LB. Inoculum was confirmed by serial dilutions. Body weight of mice was monitored, and feces were collected at different time points after infection for measuring pathogen shedding.

### Quantification of *C. rodentium* burden

On days 4, 7, 8, 12, 21, and 28 of infection, fresh fecal samples were collected, diluted in 1 mL PBS and homogenized by bead-beating with 1 mm ceramic beads once for 40 s 6000 rpm using a MagNA Lyser (Roche Diagnostics GmbH). After 7 days of *C. rodentium* infection, mice were euthanized and the colon tissue and cecal content samples were collected. Colons were harvested and colon length was measured. Colons were cut open longitudinally then cut perpendicularly into 5 pieces to be used for different assays. They were, from the distal to proximal portion assigned to histology, CFU, mRNA, cytokines, and MPO assay. Colon content was flushed out by washing with sterile PBS. Spleen and cecum samples were also harvested. Cecum and colon tissue samples were weighed and homogenized mechanically using a homogenizer (Polytron PT 2100) in PBS. To determine CFUs, serial dilutions of homogenized samples were plated on MacConkey agar plates with 100 μg/ml chloramphenicol. Plates were cultured at 37°C overnight before counting. *C. rodentium* were identified by their unique colony morphology and CFUs were calculated after normalization to the weight of each content or tissue sample.

### Constipation and gut motility measurement

A functional study of the gastrointestinal transit was performed by evaluation of water content in feces and number of fecal pellets expulsed as previously described (Stapleton et al., 2023). Briefly, on day 5 or 7 post infection, G2019S and WT mice were placed individually in cages without bedding, food, or water. The first 2 pellets of feces were collected into 1.5 ml tubes previously weighed on a precision scale. Then, tubes with sample were weighed again before and after drying and placed in a hood at room temperature, overnight. This way the degree of fecal water content could be determined. To determine the gut motility, the number of stools excreted by each animal in a time interval of 2 hours was also recorded.

### Histopathological analysis

Colon samples were obtained after euthanasia and fixed in buffered 10% formalin. Before embedding, the tissue was dehydrated in ethanol, diaphanized in xylene and embedded in paraffin. The blocks were cut using a microtome, the tissues on the slides were stained with hematoxylin and eosin (HE). Slides were scanned at a resolution of ×20 magnification, and pictures were taken using a Leica Aperio slide scanner (Leica). Histopathological scoring was conducted blindly by an expert veterinary pathologist based on the scoring criteria of colon lesions (Santos et al., 2024). They were scored based on inflammatory infiltrate (0-4), polymorphonuclear (PMN) cell infiltrate (0-4), loss of crypts (0-2), proportional loss of goblet cells (0-2), edema (0-1), erosion or ulceration (0-3), hemorrhage (0-2), and necrosis (0-1).

### Myeloperoxidase (MPO) assay

The extent of neutrophil accumulation in the colon was assessed by assaying myeloperoxidase (MPO) activity as previously described (Huang et al., 2016; Souza et al., 2000). Briefly, samples of the proximal colon were obtained after euthanasia and mechanically homogenized in a buffer (0.1 M NaCl, 0.02 M NaPO_4_, 0.015 M NaEDTA – pH 4.7) using a homogenizer (Polytron PT 2100). Samples were then centrifuged at 260 x *g* for 10 min and the pellet went through hypotonic lysis with the 0.2% NaCl solution followed by the addition of NaCl 1.6% and glucose 5% solution. Samples were centrifuged again and resuspended in 0.05 M NaPO_4_ buffer (pH 5.4) containing 0.5% hexadecyltrimethylammonium bromide and re-homogenized. Then, samples underwent to three freeze–thaw cycles using liquid nitrogen, centrifuged and supernatants were collected for the assay. The activity of MPO enzyme was measured by the quantification of changes in O.D. at 450 nm using tetramethylbenzidine (1.6 mM) and H_2_O_2_ (0.5 mM). Results were expressed as changes in absorbance O.D. of the diluted sample per g of colon tissue.

### Flow cytometry

After euthanasia, whole colons were harvested and placed in 50 ml tubes containing 20 ml of PBS with 2mM EDTA (Invitrogen, 15575-038). Tissue samples were digested with collagenase (from *Clostridium histolyticum*, Sigma, C5138) and DNAse I (grade II, Roche, 10104159001,) and colonic lamina propria (LP) immune cells were isolated (Recinto et al., 2024). Cells were stained with 1:2000 of Zombie red fixable viability dye (Biolegend, Cat#423109) in PBS for 20 min on ice. Following, Fc-Block (anti-mouse CD16/CD32) 1:100 was added (Invitrogen, Cat# 14-01061-86) and staining of surface markers with a mixture of conjugated antibodies 1:250 in 2%FBS PBS 1mM ethylenediaminetetraacetic acid (EDTA) buffer with 10% Brilliant Stain Buffer Plus (BD, 566385) for 30 min on ice. For transcriptional factor (TF) staining, cells were fixed in Foxp3/Transcription Factor Staining Buffer (eBiosciences, 00-5523-00) for 45 min on ice, then washed in permeabilization buffer and stained with a mixture of conjugated antibodies diluted 1:125 in the permeabilization buffer (listed in table 1). Single stained cells of the appropriate processed tissue were used as a reference control for unmixing. TF UltraComp eBeads (Invitrogen, 01-2222-42) were used for a single stain control of transcriptional factors and low expressed markers. Autofluorescence was deducted using unstained LP cells. Samples and their respective controls were acquired using spectral cytometer Aurora 4L (Cytek). For spectral cytometry unmixed in SpectroFlo FCS 3.0 files exported and analyzed using FlowJo v10.9 Software (BD).

**Table 1.**
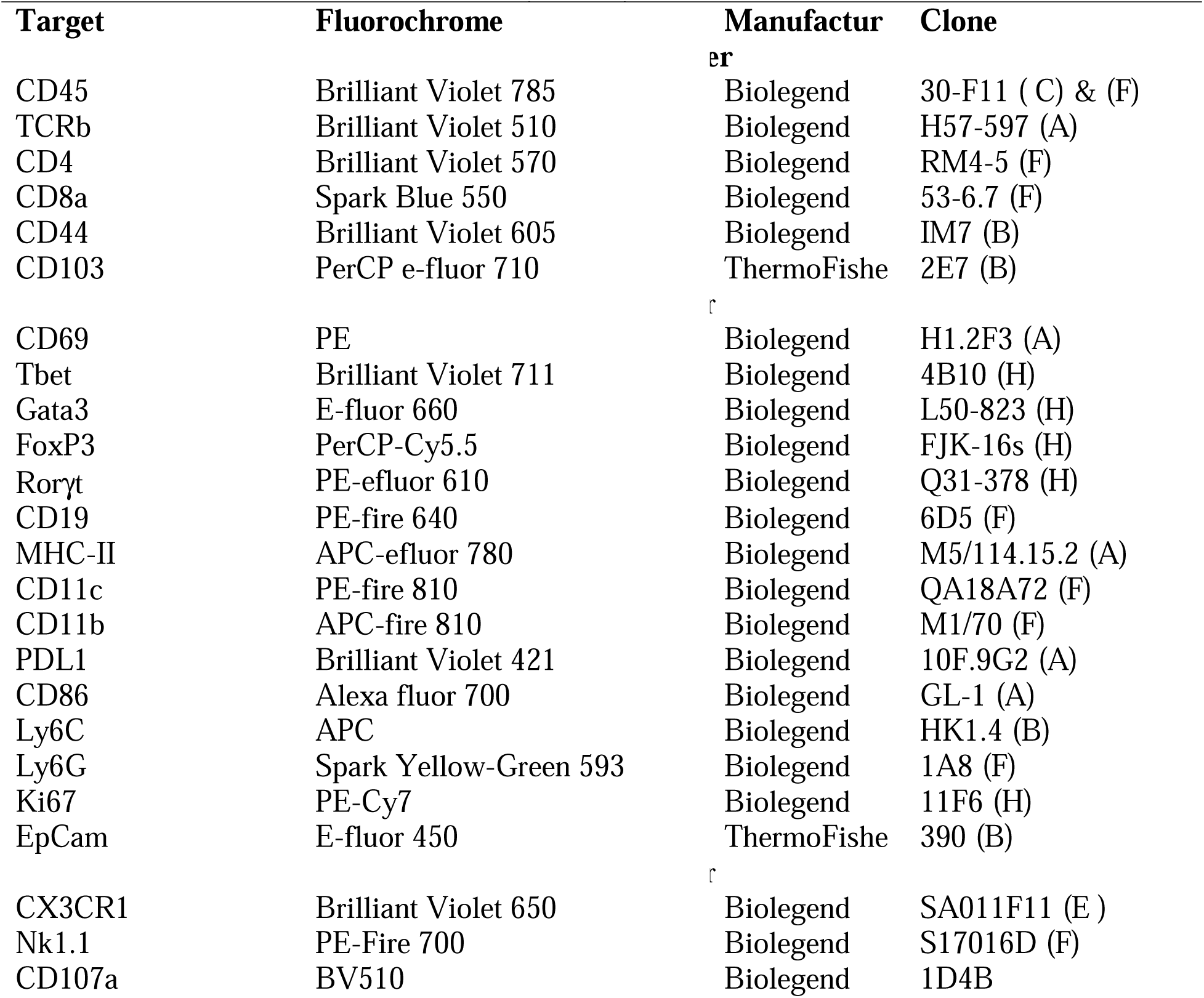
List of antibodies used in flow cytometry.

### Single cell RNA sequencing

scRNAseq was completed using 10x Genomics Chromium sequencing. Colonic lamina propria cells were isolated and pooled from three mice per condition (WT uninfected, WT infected, G2019S uninfected, G2019S infected). Using the Single Cell 3’ Reagent kit (V3.1 assay, https://www.10xgenomics.com/support/single-cell-geneexpression), 40 000 cells were loaded on the Chromium per the manufacturer’s instructions. Reverse transcription (RT), cDNA synthesis and amplification, and library preparation were completed as previously described (Recinto et al., 2024). Briefly, cells were first partitioned on a nanoliter-scale using barcoded Gel Bead-In-Emulsions (GEMs). After the GEM generation, Gel 39 Beads were dissolved, primers were released, and co-partitioned cells were lysed. Then, cellular transcripts were reverse transcribed with primers containing (1) TruSeq sequence, (2) 16 nt 10x Barcode, (3) a 12nt unique molecular identifier (UMI), and (4) a 30 nt poly(dT) sequence, which became mixed with the cell lysate and Master Mix containing RT reagents. The incubation resulted in barcoded, full-length cDNA from poly-adenylated mRNA. After that, Silane magnetic beads were used to purify the cDNA from the RT reaction mixture, followed by the amplification of cDNA through PCR, creating a library. Sequencing was performed using NovaSeq 6000 S4 PE 100bp, resulting in a final output of on average, 20 000 reads/cell, which were then processed using 10X Genomics Cell Ranger. FASTQs outputs were aligned to the mouse GRCm38.p5 reference genome. Each sample had been assigned Gene-Barcode matrices by counting UMIs and filtering non-cell associated barcodes. Finally, Seurat V4.0.6 R toolkit was used for quality control and downstream analysis of the scRNAseq data. Each Seurat object was identified with default parameters (min. cells = 3, min. features = 200) and low-quality cells and doublets were excluded based on gene counts and percent of mitochondrial genes (downstream analyses were performed on cells with gene counts between 200 and 2500, and percent of mitochondrial genes fewer than 5) (Macosko et al., 2015). Gene expression was log normalized to a scale factor of 10 000. Both Seurat objects (uninfected and infected) were then integrated as previously described in (Stuart et al., 2019).

### mRNA extraction and quantitative RT-PCR

RNA from isolated colonic immunocytes, was extracted using a Pure Link RNA kit (Invitrogen) according to the manufacturer’s instructions. RNA was reverse transcribed using SuperScript IV VILO RT (Invitrogen) as directed by the manufacturer. Quantitative PCR was performed using Taqman Fast Advanced Master Mix (Applied Biosystems). Probes are listed in Table 2. qPCR was performed on StepOnePlus (Applied Biosystems, USA). Ct values were analysed using the formula 2−ΔΔCt, normalizing target gene expression to Gapdh.

**Table 2.** List of probes used for qPCR.

| Target | Manufacturer | Cat # |
| --- | --- | --- |
| Il17a | Taqman | Mm00439618_m1 |
| Gapdh | Taqman | Mm99999915_g1 |

### Neutrophil isolation

Uninfected WT and G2019S mice were euthanized using CO_2_ affixation, femurs, tibias and peripheral blood (cardiac puncture) were harvested. Single-cell suspensions were obtained from bones by centrifugation. White blood cells from peripheral blood were isolated through repeatedly treating blood with ACK-lysis buffer to clear red blood and the total cell counts were recorded using a hemacytometer. Cells were pelleted through centrifuging at 800 × g for 10 mins at 4°C. Cells were resuspended in 900 µl/10^7^ of isolation buffer containing 0.5% Bovine Serum Albumin (BSA) and 2 mM EDTA in PBS and 10 µl/1x10^7^ of mouse anti- Ly6G MicroBeads (Miltenyi Biotec) and incubated at 4°C for 10 mins. Cells were then washed with 1 ml of isolation buffer per 1x10^7^ cells. Pelleted cells were then resuspended in 300 µl of isolation buffer and passed through magnetic-activated cell sorting (MACS) MS columns according to the manufacturer’s instructions (Miltenyi Biotec).

### *In vitro* neutrophil migration

2x10^5^ Isolated neutrophils were pelleted and resuspended in 150 µl of 37°C 10% Fetal Bovine Serum (FBS, Gibco) in Dulbecco’s Modified Eagle’s Medium (DMEM, Gibco). Neutrophils were seeded in transwell inserts with a pore size of 5 µm (VWR). Sixty microliters of mouse serum from C57BL/6 mice were added to 600 µl of 10% FBS in DMEM at the basolateral surface of the transwell. 20 µl of 1 µM N-Formylmethionine-leucyl-phenylalanine (fMLP) was supplied to the bottom well for positive control wells or 20 μl of PBS was added to the bottom well for negative control wells. For Citrobacter infections, 1 uL of bacteria (OD600 = 1) was added to 19 µl of RT sterile 1× PBS and supplied to the bottom well to obtain a multiplicity of infection (MOI) of 100. Neutrophils were given 1 h to migrate in 37°C with 5% CO2. At the end of incubation, transwells were moved to a clean 24-well plate, and the remaining liquid was aspirated. The transwell membranes were washed with 1× PBS once, residual non-migrated cells were gently removed using a cotton swab, and the membrane was fixed and stained using Kwik-Diff Stains (Epredia) as recommended by the manufacturer. Stained inserts were left to dry overnight. Membranes from dried inserts were removed by a scalpel and the output side was mounted upright onto glass slides (Fisher) using Permount Mounting Medium (Fisher Chemical). Membranes were imaged using a 10× objective on the Nikon CSU-X1 spinning disk microscope using the brightfield bypass and Nikon Digital Sight. Five random fields of view were obtained per membrane and quantified by a blinded experimenter.

### *In vitro* NETosis assay

Sterile 12mm circular coverslips (VWR) were placed in the wells of a 24-well tissue culture plate (Corning). 1x10^5^ neutrophils isolated from peripheral blood were in 1 mL of 37°C 10% Fetal Bovine Serum (FBS, Gibco) in Dulbecco’s Modified Eagle’s Medium (DMEM, Gibco) were add to the wells with coverslips. One microliter of *C. rodentium* (OD600 = 1) or 15 µl of *Pseudomonas aeruginosa* strain 6077 (OD600 = 0.45; positive control), and negative control treated cells received 15 µl of PBS. The treated cells were incubated for 2hrs at 37°C with 5% CO2. Coverslips were washed with PBS and fixed with 300 µl of 4% Paraformaldehyde (PFA, Electron Microscopy Sciences) for 30 min. The PFA was subsequently washed with PBS and the coverslips were stained with DAPI (Source) at a dilution of 1:1000 and Actin 594 (Source) at a dilution of 1:500 in 200 µl of FACS buffer for 20 min. To prepare the slides, 20 µl of ProLong Gold antifade reagent (Invitrogen) was added to glass slides (Fisher) and the coverslips were mounted that the adhered cells faced the glass slide. The glass slides were left to dry overnight. Neutrophils were imaged using the Nikon CSU-X1 spinning disk microscope. 9 fields of view from each condition were gathered and quantified using FIJI software.

### *In vitro* degranulation assay

Uninfected WT and G2019S mice were euthanized using CO_2_ and peripheral blood (cardiac puncture) with heparin were harvested. Single-cell suspensions were obtained from bones by centrifugation. White blood cells from peripheral blood were isolated through repeatedly treating blood with ACK-lysis buffer to clear red blood cells. The total cell counts were recorded using a hemacytometer. Cells were pelleted through centrifuging at 800 × g for 5 mins at 4°C. Cells were resuspended in 100 µl/10^7^ of isolation buffer containing 2% FBS and 1 mM EDTA in PBS. Neutrophils were isolated using the Ly6G-PE positive selection kit according to the manufacturer’s instructions (STEMCELL technologies, Catalog#17666). Degranulation assay was performed based on (Lorenzo-Herrero et al., 2019). Briefly, neutrophils were incubated in complete RPMI 10% FBS with or without PMA (300 ng/ml) with 4 μl /ml of anti-Lamp1 BV510 antibody for 2 h at 37°C under 5% (v/v) CO_2_. Unstimulated samples were used to detect spontaneous degranulation. After that, samples were washed with PBS acquired in flow cytometer.

### Statistical analysis

The statistical analysis was performed using GraphPad Prism version 10.2.2 (341) (Dogmatics). All data are presented as the mean ± SD and were analyzed using Two-way analysis of variance (ANOVA) followed by Tukey or Šídák post-test to compare different groups. Student’s t test was used to compare two groups. P-value<0.05 was considered a significant difference and marked as *; p<0.01 - **, p<0.001 as ***, and p<0.0001 as ****. Each experiment was done at least twice with a minimum of three mice per group.

### Illustrations

All the experimental designs and graphical illustrations were created with BioRender.com.

## Results

### LRRK2 G2019S mutation does not affect *C. rodentium* colonization or clearance

The natural murine attaching and effacing pathogen *C. rodentium* has been extensively used as a model for inducing self-limiting infectious colitis in mice, to probe intestinal inflammatory and immune responses (Connolly et al., 2018; Silberger et al., 2017; Sweet et al., 2022; Zhao et al., 2023). We sought to investigate whether PD-associated variant, LRRK2 G2019S, influenced the host response to this natural mouse pathogen. To gain insight into the role of LRRK2 G2019S in the control of *C. rodentium* infection, LRRK2 G2019S and WT mice were infected with *C. rodentium* in a single gavage with approximately 1x10^9^ CFU. Bacterial colonization was evaluated at specific time points following the infection using fecal *C. rodentium* burden as a surrogate readout (Fig.1A). By day 4 of infection, mice already presented with high levels of *C. rodentium*, indicating that the infection was well established in both WT and LRRK2 G2019S mice. By day 8 of infection, mice reached the peak of bacterial load, which decreased slightly by day 12 of infection. On day 21, bacterial clearance was progressing in both genotypes, with the majority of mice reaching the limit of CFU detection (LOD). On day 28, all the mice cleared the infection. Overall, no differences in bacterial colonization or clearance were observed between WT and LRRK2 G2019S mice during the infection (Fig. 1B). Since previous work has indicated a sex difference in some LRRK2-associated phenotypes, we also accessed larger cohorts of mice for *C. rodentium* loads at day 7 of infection, the peak of bacterial load, and compared sex differences in bacterial clearance. Female and male mice from both genotypes had similar burden of *C. rodentium* on day 7 of infection (Fig.1C).

**Figure 1.**
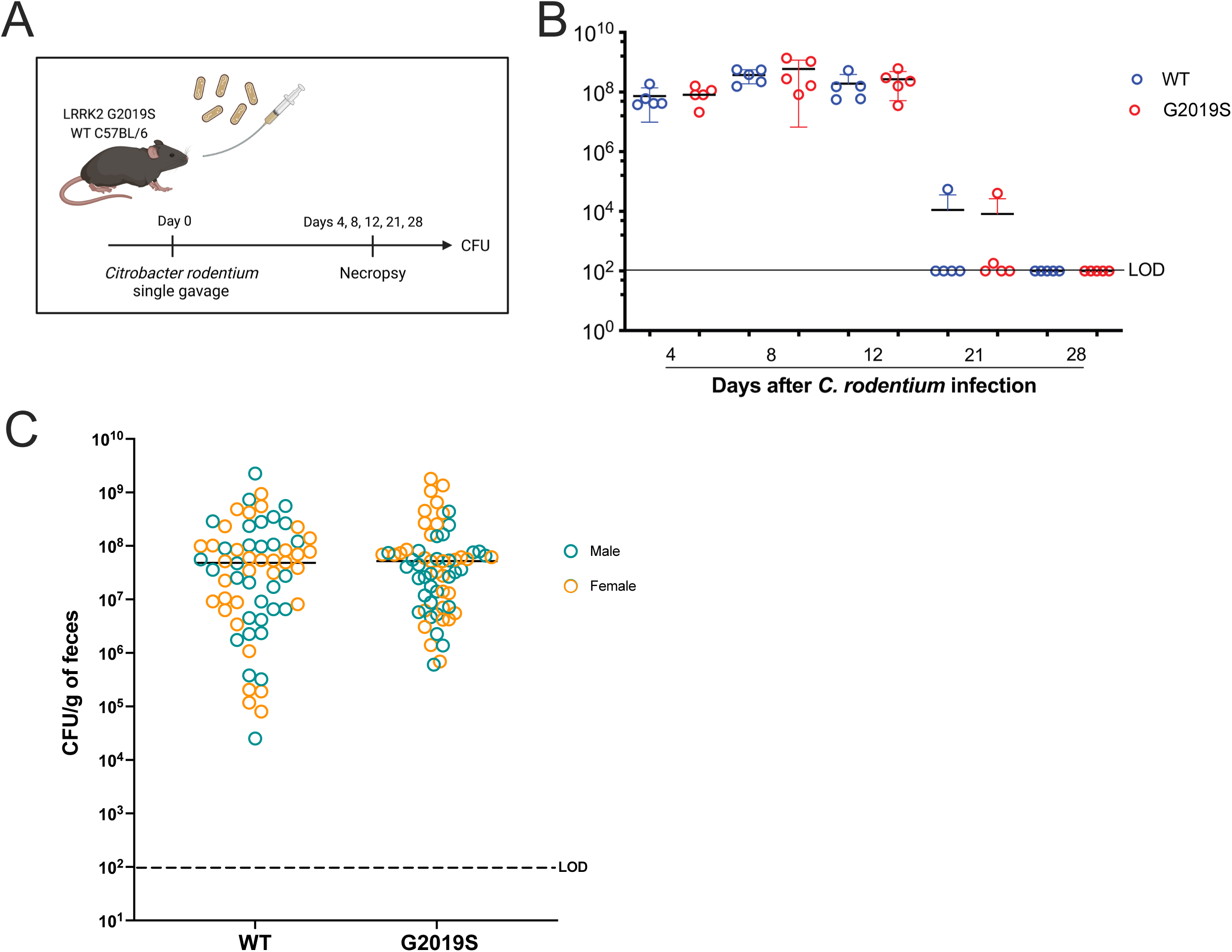
LRRK2 G2019S mutation does not interfere with *C. rodentium* colonization and clearance. (A) experimental design. Male and female LRRK2 G2019S and WT mice were gavaged once with approximately 1x10^9^ Colony Forming Units (CFUs) of *C. rodentium*. Bacterial fecal shedding was quantified on days 4, 7 or 8, 12, 21 and 28 after infection. (B) CFUs per g of feces. Data are represented as mean ± SD and analyzed by two-way ANOVA with Sidak post-test. n=5 mice per group. (C) Comparison fecal CFU *C. rodentium* burden in WT and G2019S mice on day 7 of infection with a sex separation within groups. Data are represented as mean ± SD and analyzed by t-test. Ten independent experiments are presented as a pool. n=64- 65 mice per group. Dashed lines indicate the limit of detection (LOD).

To further explore the impact of the G2019S mutation in LRRK2 during *C. rodentium* infection, we evaluated the overall response of these mice in early infection. We found that mice weight variation was similar among uninfected and infected groups and between different genotypes (Supp. Fig. 1A). Gut motility was also evaluated, by counting the number of fecal pellets excreted per hour, after 5 days of infection. Although there is a trend towards increased fecal pellets per hour after infection of WT mice, no differences were observed between the genotypes of infected mice (Supp. Fig. 1B). Moreover, on day 7 of infection, fecal water content was similar in uninfected groups and tended to decrease after infection, reaching statistical significance only for G2019S mice (Supp. Fig. 1C). Thus, indicating that LRRK2 G2019S mice have similar control of *C. rodentium* infection, with minor changes in fecal water composition.

### Single cell RNA sequencing (scRNAseq) and flow cytometry show increased neutrophil presence in LRRK2 G2019S mice following infection

While LRRK2 is known to be highly expressed in immune cells upon stimulation (Cook et al., 2017), its role in regulating inflammation and infection in the intestine remains unclear (Dikovskaya et al., 2024; Wallings & Tansey, 2019) To understand how LRRK2 G2019S mutation affects the immunophenotype of the colonic lamina propria at baseline and during infection, we completed scRNAseq. Colons were harvested from WT and LRRK2 G2019S mice at 7 days post infection, a timepoint when innate immune changes are prominent in response to *C. rodentium* infection (Mullineaux-Sanders et al., 2019) In parallel, colons were also collected from age-and sex-matched uninfected WT and LRRK2 G2019S controls. Using *Seurat*, we integrated datasets from our four conditions, following standard quality control metrics (number of unique molecular identifiers (UMIs) per cell, percent mitochondrial reads) (Stuart et al., 2019). Post filtering, we yielded a total of 13,892 cells sequenced at a depth of ∼20,000 genes per cell, depicted on respective UMAP plots (Fig. 2A). Through unsupervised clustering and manual annotation based on established transcriptional markers for immunocytes within the lamina propria, all major immune cell types were identified (Blecher-Gonen et al., 2019; Recinto et al., 2024) (Fig. 2B). Although at baseline there were no noticeable differences in cluster sizes between genotypes, the data from infected mice indicated increased representation of neutrophils in the LRRK2 G2019S sample compared to WT sample (Fig. 2A). Between the four conditions, *Lrrk2* expression was low to moderate (4-8% of total immunocytes) which is consistent with other published datasets, showing that detection of *Lrrk2* mRNA and protein can be challenging (Biskup et al., 2007; Gardet et al., 2010). Nonetheless, LRRK2 G2019S-infected immunocytes showed higher average *Lrrk2* expression relative to all other conditions (Fig. 2C).

**Figure 2.**
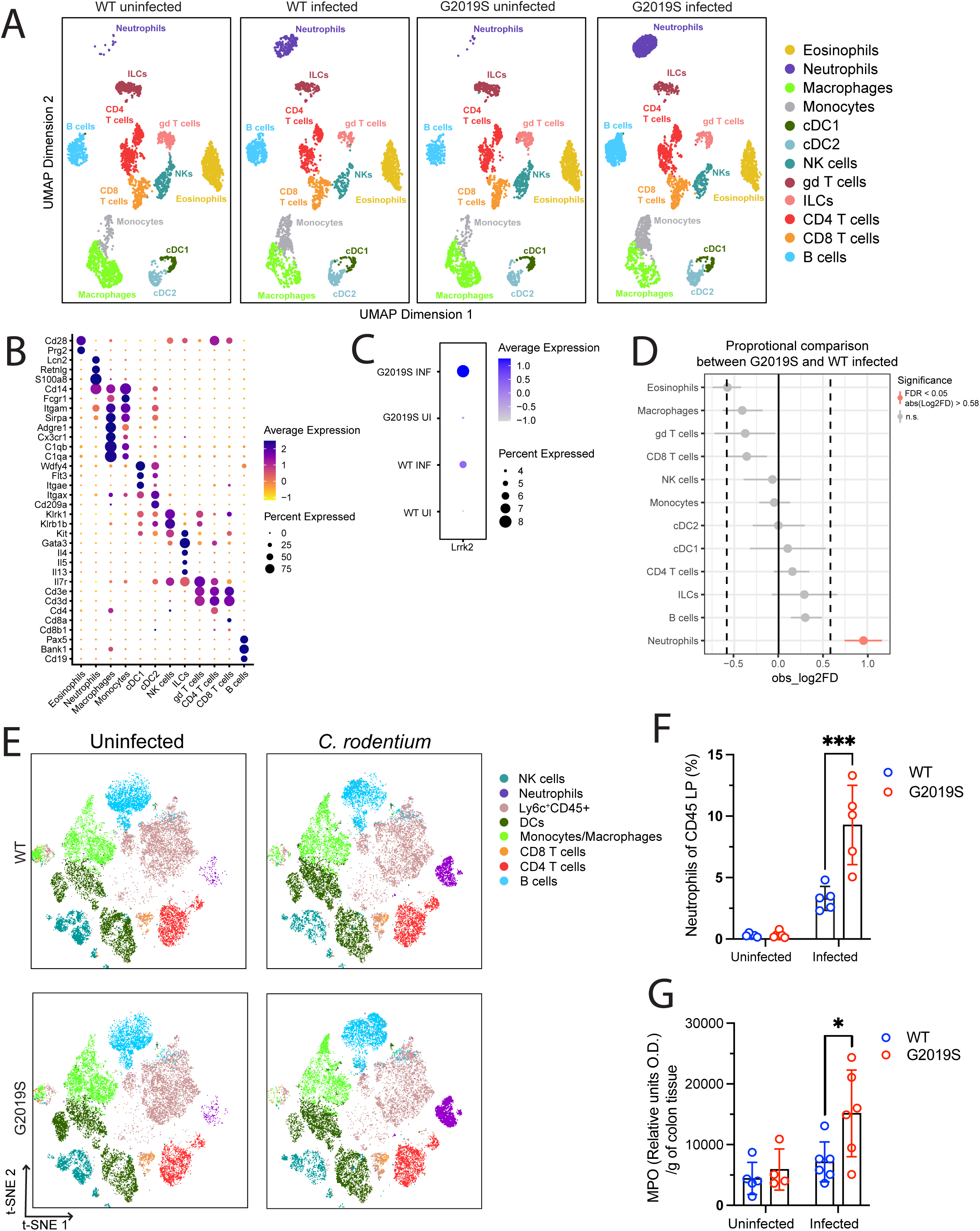
Single cell RNA sequencing (scRNAseq) and flow cytometry show increased neutrophil presence in colonic lamina propria of LRRK2 G2019S mice following infection. (A) Unsupervised clustering of immune cells in WT or G2019S uninfected and infected mice. The scRNAseq datatset contains 13,892 cells total sequenced at a depth of ∼20,000 genes per cell, depicted on respective UMAP plots. scRNAseq obtained from pooled conditions of n = 3 mice per group (B) Dotplot depicting identifying markers used to annotate each immune cluster based off literature (C) Dotplot depicting average *Lrrk2* expression across all immune cell clusters, split by condition. (D) Proportional analysis conducted through permutation test comparing G2019S infected and WT infected conditions. Significance cut-off: FDR < 0.05. (E) Flow cytometry results of immune cell cluster projected onto t-SNE plots. (F) Percent Ly6G- positive neutrophils isolated from CD45+ immune cells of the colonic lamina propria. Data are represented as mean ± SD and analyzed by two-way ANOVA with Sidak post-test. n=4-5 mice per group. * < 0.05 and *** p < 0.001. One independent experiment is presented. (G) MPO enzymatic assay results of colonic tissue sample measured through O.D. and normalized over gram of colonic tissue. Data are represented as mean ± SD and analyzed by two-way ANOVA with Sidak post-test. n=4-6 mice per group. * p < 0.05 one representative of two independent experiments are presented.

To comprehend how LRRK2 G2019S activity may alter immunocyte proportions in the lamina propria under infectious conditions, we completed a proportional analysis of LRRK2 G2019S against WT infection. Confirming what was observed in Figure 2A, neutrophils were the only cluster with a significant proportional upregulation in LRRK2 G2019S infected mice (Fig. 2D). Next, to validate our scRNAseq findings at a protein level, we immunophenotyped the colonic lamina propria of WT and LRRK2 G2019S mice at baseline and at day 7 post infection through flow cytometry (Fig. 2E and Table 1) t-SNE pseudo dot plots of cells in lamina propria showed that while there were no distinguishable genotype differences between cluster sizes in uninfected mice, there was a significant increase in neutrophils in LRRK2 G2019S infected mice, compared to WT (Fig. 2E-F). The increase in neutrophils was confirmed by the activity of myeloperoxidase (MPO), an indirect measure of neutrophil presence (Hanning et al., 2023). As expected, LRRK2 G2019S infected mice showed higher levels of MPO activity compared to WT infected (Fig. 2G). Together, our findings demonstrate a marked increase in neutrophils in the colonic lamina propria of LRRK2 G2019S infected mice, at mRNA and protein levels.

### Differentially expressed gene analysis indicates neutrophils of LRRK2 G2019S infected mice have an increased pro-inflammatory profile

To understand how LRRK2 G2019S mutation affects gene regulation, differential gene expression analysis was performed on each immune cell cluster, comparing WT and LRRK2 G2019S-infected mice. The number of significant up and downregulated gene hits for each cluster are represented in a heatmap (Fig. 3A). Among all the clusters, neutrophils, monocytes, and γδ T cells had the highest number of differentially expressed genes (DEGs). In LRRK2 G2019S infected neutrophils, there were 22 significantly upregulated genes, and 9 significantly downregulated genes, compared to WT mice (Table 3 and 4). Several upregulated genes of interest include neutrophil migration and maturation markers *Plaur, Fgl2,* the gamma-actin encoding gene *Actg1,* chemokines *Cxcl10, Ccl6, Ccr1*, and interferon (IFN) signaling-related genes *Irf1, Irgm1,* and *Stat1* (Gyetko et al., 1995). Using our total differentially regulated gene list, we conducted pathway enrichment analysis through Gene Ontological (GO) terms to identify key associated biological pathways. GO terms pointed to pro-inflammatory signaling, including increased IFN-mediated signaling (type II IFN-mediated signaling, defense/response to virus, and cytokine-mediate signaling), cellular response to IFN-β, and cytolysis disruption of another organism (defense response to bacterium and cell killing) (Fig. 3B).

**Figure 3.**
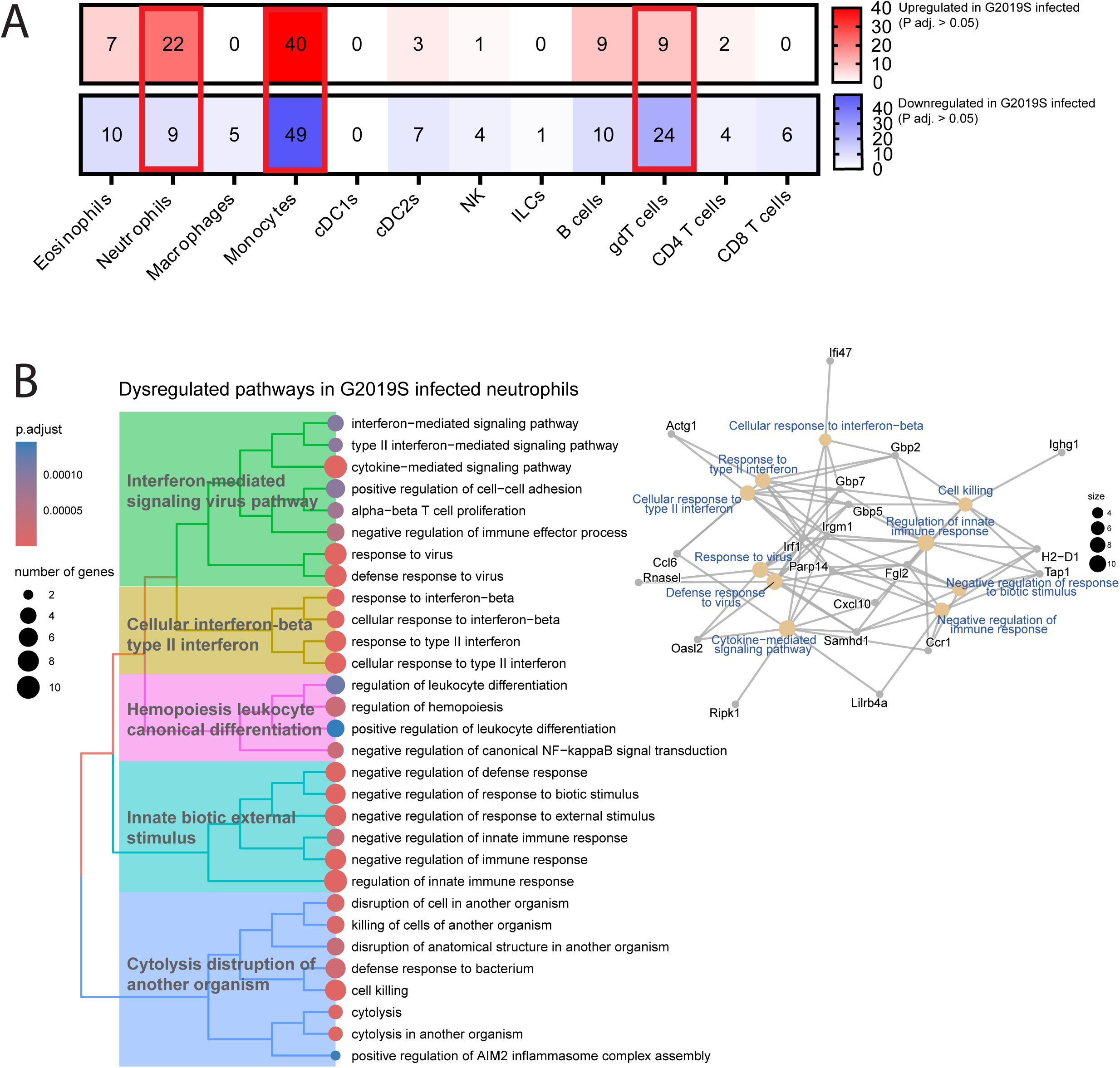
Differentially expressed gene analysis indicates neutrophils of LRRK2 G2019S infected mice have an increased pro-inflammatory profile. (A) Differential gene expression (DEG) analysis comparing G2019S and WT infected mice for each immune cell cluster calculated using Seurat differential expression analysis by non-parametric Wilcoxon rank sum test with Bonferroni correction. Adjusted P value cut off = 0.05, Q value cut off = 0.2. Upregulated genes depicted as a heatmap in red and downregulated depicted in blue. (B) Pathway enrichment analysis completed on DEGs obtained by comparing G2019S and WT infected neutrophils. DEGs were run against Gene ontology (GO) biological pathways database using *ClusterProfiler.* P value cut off = 0.05, q value cut off = 0.2. Top 30 GO term categories depicted as a Treeplot (Left). Cnetplots depict association of differentially expressed gene hits, and which GO term pathways they correspond to (Right).

**Table 3.** Top 20 significantly upregulated differentially expressed genes in G2019S infected neutrophils compared to WT infected.

| Genes | p_val | avg_log2FC | pct.1 | pct.2 | p_val_adj |
| --- | --- | --- | --- | --- | --- |
| Plaur | 4.2896E-17 | 2.56646846 | 0.722 | 0.466 | 8.9249E-13 |
| Irf1 | 3.224E-13 | 2.9665498 | 0.79 | 0.561 | 6.7079E-09 |
| Actg1 | 1.9157E-10 | 17.3159898 | 0.982 | 0.951 | 3.9858E-06 |
| Fgl2 | 8.2326E-10 | 9.70633498 | 0.757 | 0.587 | 1.7129E-05 |
| Tap1 | 1.5713E-09 | 1.59248386 | 0.572 | 0.323 | 3.2693E-05 |
| Irgm1 | 1.6212E-09 | 5.40865353 | 0.489 | 0.256 | 3.3731E-05 |
| Cxcl10 | 2.6077E-09 | 60.5959953 | 0.491 | 0.269 | 5.4256E-05 |
| Samhd1 | 1.4522E-08 | 11.5421141 | 0.888 | 0.776 | 0.00030215 |
| Rhoh | 6.3858E-08 | 2.65791361 | 0.402 | 0.202 | 0.00132863 |
| Ccl6 | 2.1192E-07 | 14.5666966 | 0.683 | 0.48 | 0.00440919 |
| Gbp7 | 2.6187E-07 | 3.02910595 | 0.527 | 0.318 | 0.00544852 |
| Gbp5 | 4.5256E-07 | 15.8374442 | 0.593 | 0.408 | 0.00941592 |
| Rnasel | 5.4518E-07 | 8.45985251 | 0.375 | 0.188 | 0.01134309 |
| Ccr1 | 6.2266E-07 | 3.79811961 | 0.86 | 0.74 | 0.01295515 |
| Nampt | 6.7531E-07 | 2.56407938 | 0.579 | 0.381 | 0.01405041 |
| Snx20 | 7.6411E-07 | 6.43647727 | 0.902 | 0.865 | 0.01589812 |
| Srgn | 9.5358E-07 | 40.3982648 | 0.998 | 1 | 0.01984016 |
| H2-D1 | 1.0444E-06 | 3.58743132 | 0.985 | 0.951 | 0.02173024 |
| Gdap10 | 1.3744E-06 | 2.6104548 | 0.36 | 0.184 | 0.02859656 |
| Ifi47 | 1.5055E-06 | 7.95125562 | 0.428 | 0.238 | 0.03132359 |

**Table 4.**
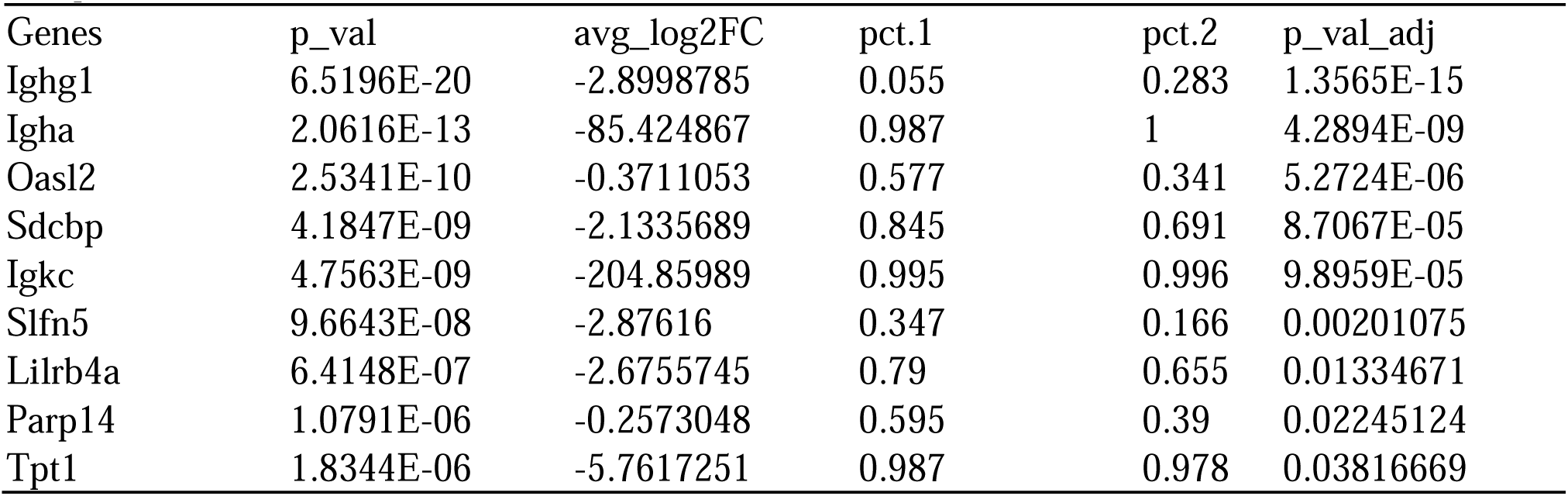
Top downregulated differentially expressed genes in G2019S infected neutrophils compared to WT infected.

| Genes | p_val | avg_log2FC | pct.1 | pct.2 | p_val_adj |
| --- | --- | --- | --- | --- | --- |
| Ighg1 | 6.5196E-20 | -2.8998785 | 0.055 | 0.283 | 1.3565E-15 |
| Igha | 2.0616E-13 | -85.424867 | 0.987 | 1 | 4.2894E-09 |
| Oasl2 | 2.5341E-10 | -0.3711053 | 0.577 | 0.341 | 5.2724E-06 |
| Sdcbp | 4.1847E-09 | -2.1335689 | 0.845 | 0.691 | 8.7067E-05 |
| Igkc | 4.7563E-09 | -204.85989 | 0.995 | 0.996 | 9.8959E-05 |
| Slfn5 | 9.6643E-08 | -2.87616 | 0.347 | 0.166 | 0.00201075 |
| Lilrb4a | 6.4148E-07 | -2.6755745 | 0.79 | 0.655 | 0.01334671 |
| Parp14 | 1.0791E-06 | -0.2573048 | 0.595 | 0.39 | 0.02245124 |
| Tpt1 | 1.8344E-06 | -5.7617251 | 0.987 | 0.978 | 0.03816669 |

LRRK2 G2019S infected monocytes showed 40 upregulated genes and 49 downregulated genes, of which the top twenty are depicted in Table 5 and 6. GO term analysis indicated dysregulation in cellular response to infection and presentation of peptides via MHC II (Supp. Fig. 2A). Specifically, changes in MHC II peptide presentation were driven by downregulation of genes – *H2-Aa* and *H2-Ab1* which encode α and β components of the MHC II complex, respectively. In addition to gene expression differences, flow cytometry revealed increased influx of Ly6c^High^ monocytes with decreased MHC II expression in LRRK2 G2019S infected mice (Supp. Fig. 2B, data not shown). This was further supported through sub-clustering of monocytes in the scRNAseq dataset. Monocytes were sub-clustered into Ly6c2^+^, Cxcr1^+^, or Itgax^+^, indicating a more immature, macrophage-like, or DC-like profile, respectively (Supp. Fig. 2C). Proportional permutation analysis showed that Ly6C^high^-monocytes were significantly upregulated in the G2019S infected condition compared to WT infected, supporting our flow cytometry data (Supp. Fig. 2D). LRRK2 G2019S infected γδ T cells showed 9 upregulated genes and 24 downregulated genes, depicted in Table 7 and 8. Using these genes, GO term analysis was completed and indicated that most hits were associated with ribosomal processes (ribosome biogenesis, rRNA processing, rRNA homeostasis) (Supp. Fig. 3A). Taking these results together, we can determine that the LRRK2 G2019S mutation affects the immune compartment during infection, leading to dysregulated gene expression in neutrophils, monocytes, and γδ T cells.

**Table 5.** Top 20 upregulated differentially expressed genes in G2019S infected monocytes compared to WT infected.

| Genes | p_val | avg_log2FC | pct.1 | pct.2 | p_val_adj |
| --- | --- | --- | --- | --- | --- |
| Igha | 1.0571E-29 | 25.3474125 | 0.994 | 1 | 2.1993E-25 |
| Cd74 | 1.755E-12 | 21.20771 | 0.901 | 0.962 | 3.6515E-08 |
| Tspo | 1.8415E-12 | 4.19838292 | 0.968 | 0.88 | 3.8314E-08 |
| Rpl23 | 4.0434E-12 | 24.0834363 | 0.996 | 1 | 8.4127E-08 |
| Rps8 | 5.4179E-12 | 21.136032 | 1 | 1 | 1.1272E-07 |
| Rpl37 | 1.1689E-11 | 12.9243146 | 0.998 | 1 | 2.4319E-07 |
| Gbp2 | 2.0159E-11 | 21.2339402 | 0.749 | 0.609 | 4.1942E-07 |
| Rplp1 | 3.1311E-11 | 9.63968894 | 1 | 1 | 6.5145E-07 |
| H2-Eb1 | 4.5007E-11 | 11.1088873 | 0.692 | 0.833 | 9.3641E-07 |
| Rps26 | 1.2657E-10 | 1.00843319 | 0.994 | 0.997 | 2.6333E-06 |
| Psme2 | 1.9543E-10 | 9.32780777 | 0.872 | 0.814 | 4.0661E-06 |
| Apoe | 8.4277E-10 | 112.097545 | 0.919 | 0.915 | 1.7535E-05 |
| Rpl30 | 2.1667E-09 | 7.56304984 | 0.996 | 1 | 4.508E-05 |
| Rpl26 | 3E-09 | 22.4384066 | 0.996 | 0.995 | 6.2419E-05 |
| AA467197 | 4.4218E-09 | 42.4054799 | 0.37 | 0.205 | 9.2E-05 |
| Prdx1 | 9.1838E-09 | 88.8311918 | 0.86 | 0.746 | 0.00019108 |
| Rpl37a | 1.0184E-08 | 18.7438149 | 1 | 1 | 0.00021188 |
| Rel | 1.2525E-08 | 9.6740115 | 0.931 | 0.967 | 0.00026059 |
| Actb | 1.2586E-08 | 38.5187837 | 1 | 1 | 0.00026186 |

**Table 6.** Top 20 downregulated differentially expressed genes in G2019S infected monocytes compared to WT infected.

| Genes | p_val | avg_log2FC | pct.1 | pct.2 | p_val_adj |
| --- | --- | --- | --- | --- | --- |
| Rpl41 | 2.6862E-22 | -18.989704 | 1 | 0.997 | 5.5888E-18 |
| Igkc | 1.5938E-18 | -32.17178 | 0.998 | 0.997 | 3.3161E-14 |
| Rpl35a | 8.2608E-17 | -20.532874 | 1 | 1 | 1.7188E-12 |
| Gm34084 | 4.1677E-15 | -5.9348725 | 0.271 | 0.511 | 8.6714E-11 |
| H2-Aa | 3.9627E-14 | -13.416923 | 0.761 | 0.877 | 8.2449E-10 |
| Fau | 9.2198E-14 | -13.41302 | 1 | 1 | 1.9183E-09 |
| Ighg1 | 6.7423E-13 | -2.0926061 | 0.087 | 0.268 | 1.4028E-08 |
| H2-Ab1 | 1.2516E-12 | -59.583164 | 0.773 | 0.877 | 2.6041E-08 |
| Rps27rt | 1.8053E-12 | -1.4087159 | 0.253 | 0.473 | 3.7561E-08 |
| Malat1 | 5.7462E-12 | -413.04345 | 1 | 1 | 1.1956E-07 |
| Ptpn18 | 6.7478E-12 | -2.2165025 | 0.787 | 0.915 | 1.4039E-07 |
| Selenop | 1.755E-11 | -13.365385 | 0.623 | 0.768 | 3.6514E-07 |
| Rhog | 2.3855E-10 | -1.4665926 | 0.721 | 0.828 | 4.9633E-06 |
| Rps15 | 2.4367E-10 | -10.240873 | 0.996 | 1 | 5.0699E-06 |
| Zbp1 | 3.2779E-10 | -1.3702754 | 0.771 | 0.546 | 6.8201E-06 |
| Rps20 | 3.4633E-10 | -1.816356 | 0.998 | 1 | 7.2057E-06 |
| Rpl34 | 3.5083E-10 | -19.576476 | 1 | 0.995 | 7.2993E-06 |
| Rps28 | 5.7063E-10 | -11.689014 | 0.998 | 1 | 1.1873E-05 |
| Rgs10 | 4.7215E-09 | -2.4752542 | 0.472 | 0.631 | 9.8236E-05 |
| Rps15a | 7.0701E-09 | -6.1423425 | 0.99 | 0.997 | 0.0001471 |

**Table 7.** Top 20 upregulated differentially expressed genes in G2019S infected gd T cells compared to WT infected.

| Genes | p_val | avg_log2FC | pct.1 | pct.2 | p_val_adj |
| --- | --- | --- | --- | --- | --- |
| Igha | 3.7823E-11 | 1.00807578 | 0.987 | 1 | 7.8696E-07 |
| Rpl35a | 7.0365E-11 | 0.26855508 | 0.987 | 1 | 1.464E-06 |
| Rps29 | 7.5225E-10 | 25.1585336 | 0.993 | 1 | 1.5651E-05 |
| Rpl6 | 4.4485E-09 | 0.96552265 | 1 | 1 | 9.2556E-05 |
| Rps2 | 2.544E-08 | 10.2471969 | 0.974 | 0.989 | 0.00052931 |
| Rpl28 | 2.2513E-07 | 1.77281773 | 0.967 | 1 | 0.00468415 |
| Rpl37 | 6.4949E-07 | 0.6131466 | 0.993 | 0.989 | 0.01351332 |
| Igkc | 6.6611E-07 | 72.2453023 | 1 | 1 | 0.01385915 |
| Pdia3 | 1.5306E-06 | 7.76599087 | 0.684 | 0.876 | 0.03184567 |

**Table 8.**
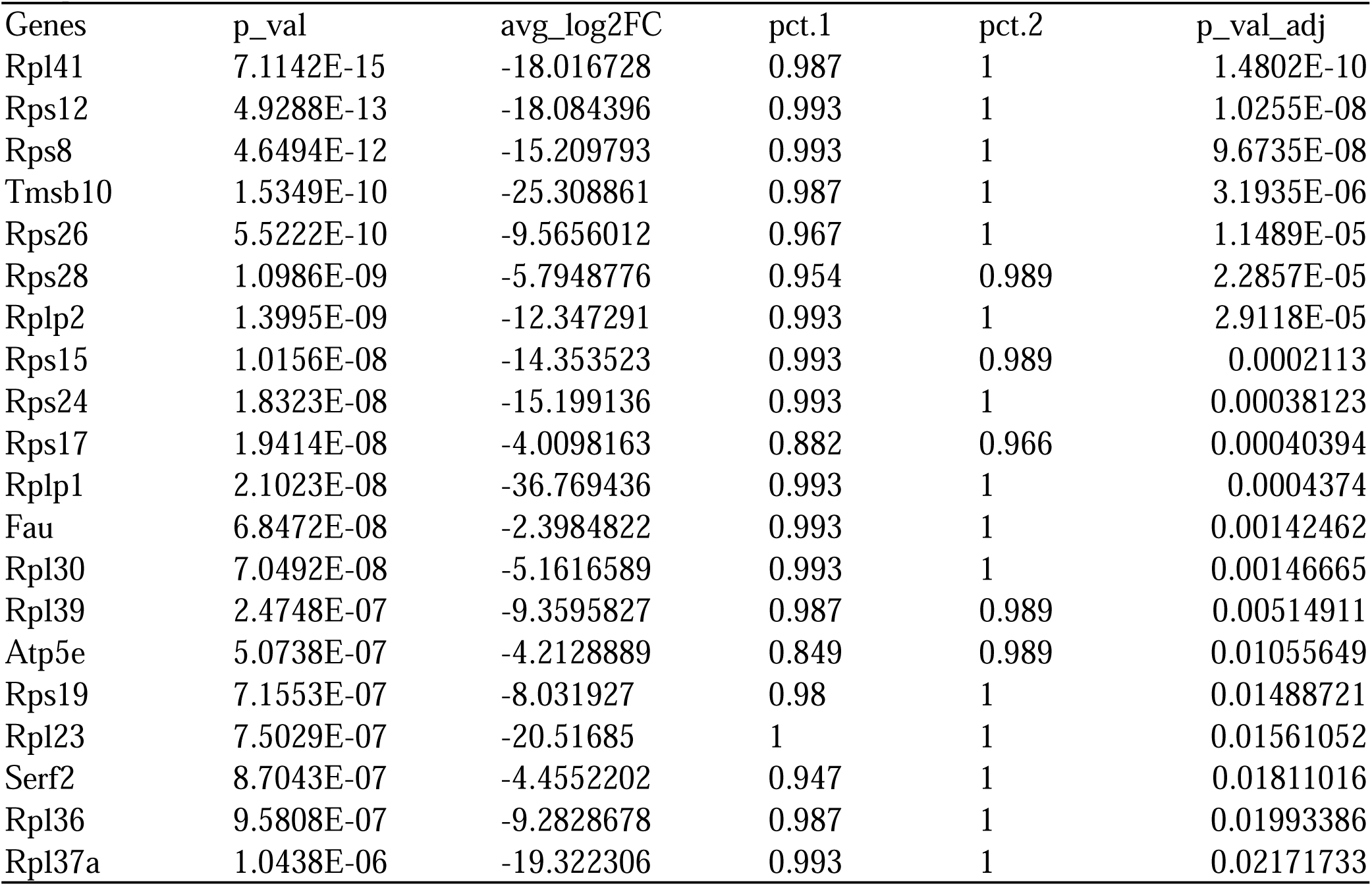
Top 20 downregulated differentially expressed genes in G2019S infected gd T cells compared to WT infected.

### LRRK2 G2019S mutation leads to increased colon morphological damage after *C. rodentium* infection

Given the differences in the innate immune response, we next evaluated the impact of the G2019S mutation at a tissue level, after infection. Therefore, we performed histopathological analysis of colonic tissue from *C. rodentium*-infected WT and G2019S LRRK2 mice on day 7 of infection. Uninfected mice showed minimal signs of disease, with no differences between genotypes. Infected mice demonstrated elevated inflammatory infiltrate (green arrows), erosion (red arrows), goblet cells loss (yellow stars), and edema (blue stars). Although those features were present in both groups, they were more evident in LRRK2 G2019S, which displayed a significant increase in the total histopathological score (Fig. 4A-B and Supp. Fig. 4A-H). Together, these results suggest that LRRK2 G2019S mice undergo a more severe inflammatory process in the colon tissue compared to WT mice during infection.

**Figure 4.**
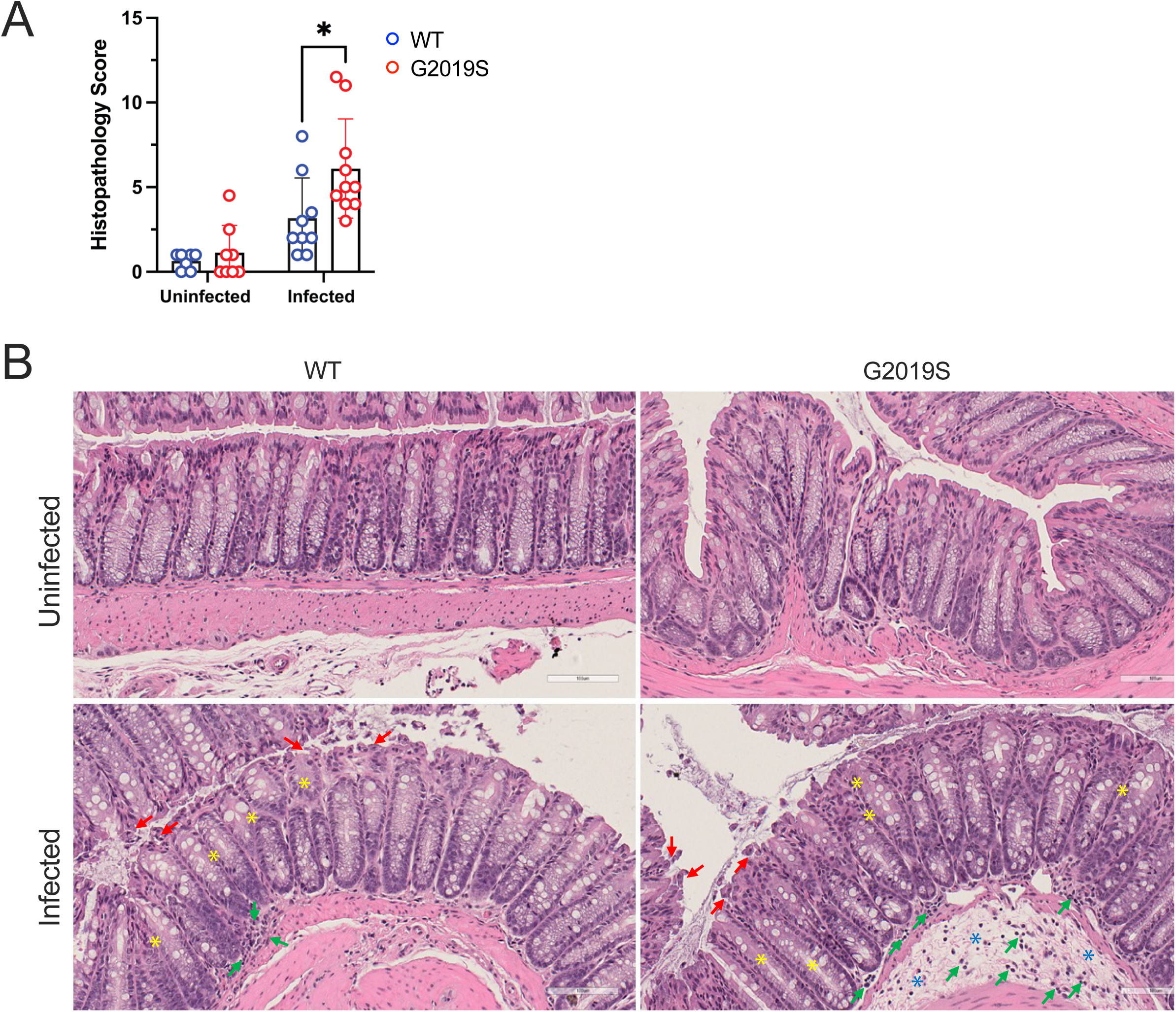
LRRK2 G2019S mutation leads to increased colon morphological damage after *C. rodentium* infection. Male and female LRRK2 G2019S and WT mice were infected once with approximately 1x10^9^ CFU of *C. rodentium*. Colons were harvested 7 days after infection, fixed in 10% formalin for 24 h, washed in 70% ethanol, and diaphanized in xylene before embedding in paraffin. Sections were stained with Hematoxylin and Eosin (H&E) and scored for histopathology. (A) Total histopathology score. (B) H&E representative images. Data are presented as mean ± SD and analyzed by two-way ANOVA with Sidak post-test. * p< 0.05. n= 7-11 mice per group. Two independent experiments are polled. White bars represent 100 µm. Symbols: blue star, edema; green arrow, inflammatory infiltrate; red arrow, erosion; yellow star, goblet cell loss. Please, also see Supplementary figure 4.

### LRRK2 G2019S mutation leads to a cell-intrinsic increase in neutrophil effector function

Based on the increased presence and dysregulated gene expression in neutrophils in infected LRRK2 G2019S mice, together with previous data indicating neutrophils as a cell type expressing high levels of LRRK2 (Taylor & Alessi, 2020), we wanted to determine if the LRRK2 G2019S mutation drove cell-intrinsic differences in neutrophils and their responses to infectious or inflammatory stimuli. Notably, several of the differentially expressed genes uncovered in the scRNAseq data have previously been implicated in neutrophil effector functions such as migration (Gyetko et al., 1995), degranulation (Skokowa et al., 2009), and NETosis (Li et al., 2022). To assess degranulation, Ly6G-purified blood neutrophils were exposed to phorbol myristate acetate (PMA) and assessed for surface expression of Lamp1 using flow cytometry. Both genotypes displayed increased degranulation upon exposure to PMA, with LRRK2 G2019S neutrophils showing significantly more degranulation than WT neutrophils (Fig. 5A). Similarly, in 1 hour migration assays using purified neutrophils (Fig. 5 B-C) significantly more LRRK2 G2019S neutrophils migrated towards both *C. rodentium* or the chemoattractant fMLP compared to WT neutrophils. NETosis was also significantly increased in LRRK2 G2019S neutrophils, in comparison to WT neutrophils, following exposure to *C. rodentium* (Fig. 5D-E). Notably, in the migration and NETosis assays with *C. rodentium* the shape of the LRRK2 G2019S neutrophils was noticeably distorted with a marked difference in actin staining present in the NETosis samples (Fig. 5D). Taken together, these results indicate that the LRRK2 G2019S mutation has a profound cell-intrinsic impact on neutrophil function, resulting in an overall increase in neutrophil effector mechanisms.

**Figure 5.**
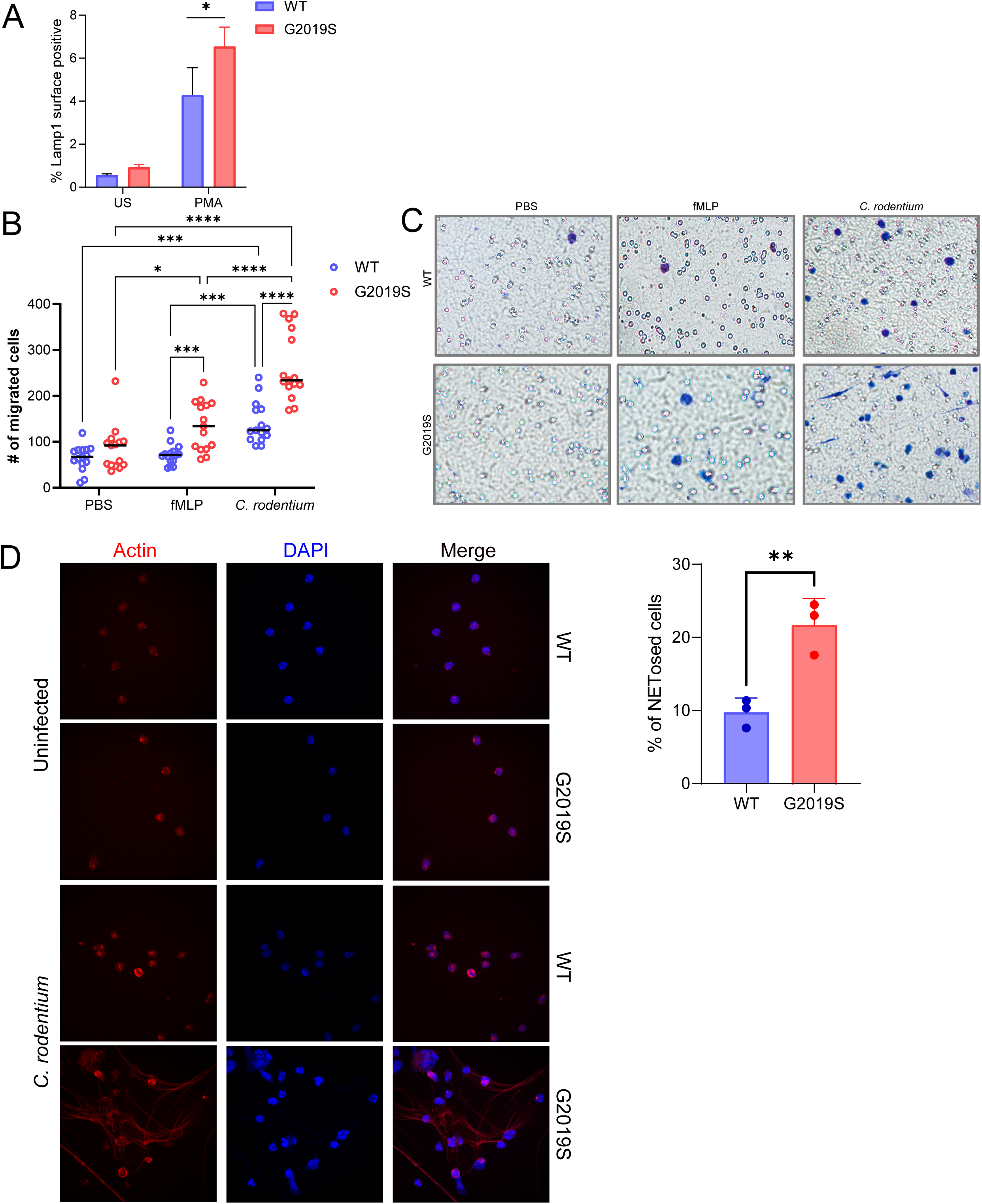
LRRK2 G2019S mutation leads to a cell-intrinsic increase in neutrophil effector function. Uninfected WT and G2019S mice were euthanized using CO_2_ affixation, femurs, tibias and peripheral blood were harvested. Red blood cells were lysed, and Ly6G-isolation was performed. (A) Blood neutrophil degranulation. Neutrophils were incubated with PMA for 2 h at 37°C with anti-LAMP-1 antibody and acquired on flow cytometer. Data are presented as mean ± SD and analyzed by two-way ANOVA with Sidak post-test. * p< 0.05. n=4 mice per group (pooled). One independent experiment is presented as a pool of mice, technical triplicates were used for quantification. (B-C) Number of migrated cells. Bone Marrow-isolated neutrophils were seeded in TransWell inserts towards fMLP or *C. rodentium* and incubated for one hour at 37°C. At the end of incubation, TransWells were moved to a clean 24-well plate, and the remaining liquid was aspirated. The TransWell membranes were washed, residual non-migrated cells were removed, and the membrane was fixed and stained. Membranes from dried inserts were mounted upright onto glass slides and imaged. Five random fields of view were obtained per membrane and were quantified by a blinded experimenter. Data are presented as mean ± SD and analyzed by two-way ANOVA with Sidak post-test. * p< 0.05, ** p<0.01, p < 0.001, p<0.0001. n=3 mice per group. One independent experiment is presented. (D-E) Blood NETosis. Blood Ly6G- isolated neutrophils 1x10^5^ neutrophils were added to the 24-well plate with coverslips. Cells were stimulated with *C. rodentium* and incubated for 2h at 37°C. Coverslips were washed with PBS and fixed. Then, stained for DAPI and actin594 and imaged. Nine random fields of view from each condition were gathered and quantified results are showed as the percentage of NETosed cells. Data are presented as mean ± SD and analyzed by two-way ANOVA with Sidak post-test. * p< 0.05, ** p<0.01, p < 0.001, p<0.0001. n=3 mice per group. One independent experiment is presented.

### Colonic infection in LRRK2 G2019S mice initiates a skewing towards Th17 CD4^+^ T cells

Recent studies have shown that neutrophils can regulate Th17 cells through the formation of neutrophil extracellular traps (NETs), which can cause tissue damage and prolonged inflammatory response (Fan et al., 2023; Wang et al., 2024). Th17 cells are also prominent players in response to *C. rodentium* infection (Li et al., 2014; Stockinger, 2021; Wang et al., 2014). We further investigated CD4+ T cell subsets. Using unsupervised clustering, we identified six distinct subsets – T naïve, Th1, Th2, Th17, and IL-10-negative and positive T regulatory cells (Treg IL10^-^, and Treg IL10^+^) in accordance with established transcriptional markers for CD4+ T cells (Stubbington et al., 2015) (Fig. 6A-B). At baseline, no genotype differences between cell clusters were identified. However, following infection, Th17 cells demonstrated a trend towards increase in LRRK2 G2019S infected condition compared to WT infected (Fig. 6A). Using flow cytometry, we also validated the tendency towards increase in LRRK2 G2019S infected mice (Fig. 6C). To better identify if there is a greater Th17 effector skew in LRRK2 G2019S infected mice, we looked at the average expression of IL-17A and IL-17F, two major cytokines produced by Th17 cells (Ouyang et al., 2008). Additionally, we assessed transcriptional factor RAR-related orphan receptor gamma (RORyt), encoded by *Rorc*, which is a critical regulator of Th17 differentiation, inducing expression of IL-17A and IL-17F (Fan et al., 2023; Ivanov et al., 2006). We saw tendencies for increase in *Il17a* expression in the Th17 cluster of LRRK2 G2019S infected mice, compared to other conditions (Fig. 6D). Reverse transcriptase quantitative PCR (RTqPCR) on CD45^+^ immunocytes from the colonic lamina propria further demonstrated a significant upregulation of *Il17a* in LRRK2 G2019S infected mice (Fig. 6E). To better understand if a skew towards Th17 is occurring, we also looked at the Th17/T naive ratio. Both through scRNAseq and flow cytometry, we saw a tendency to increase in this ratio in LRRK2 G2019S infected mice (Fig. 6F).

**Figure 6.**
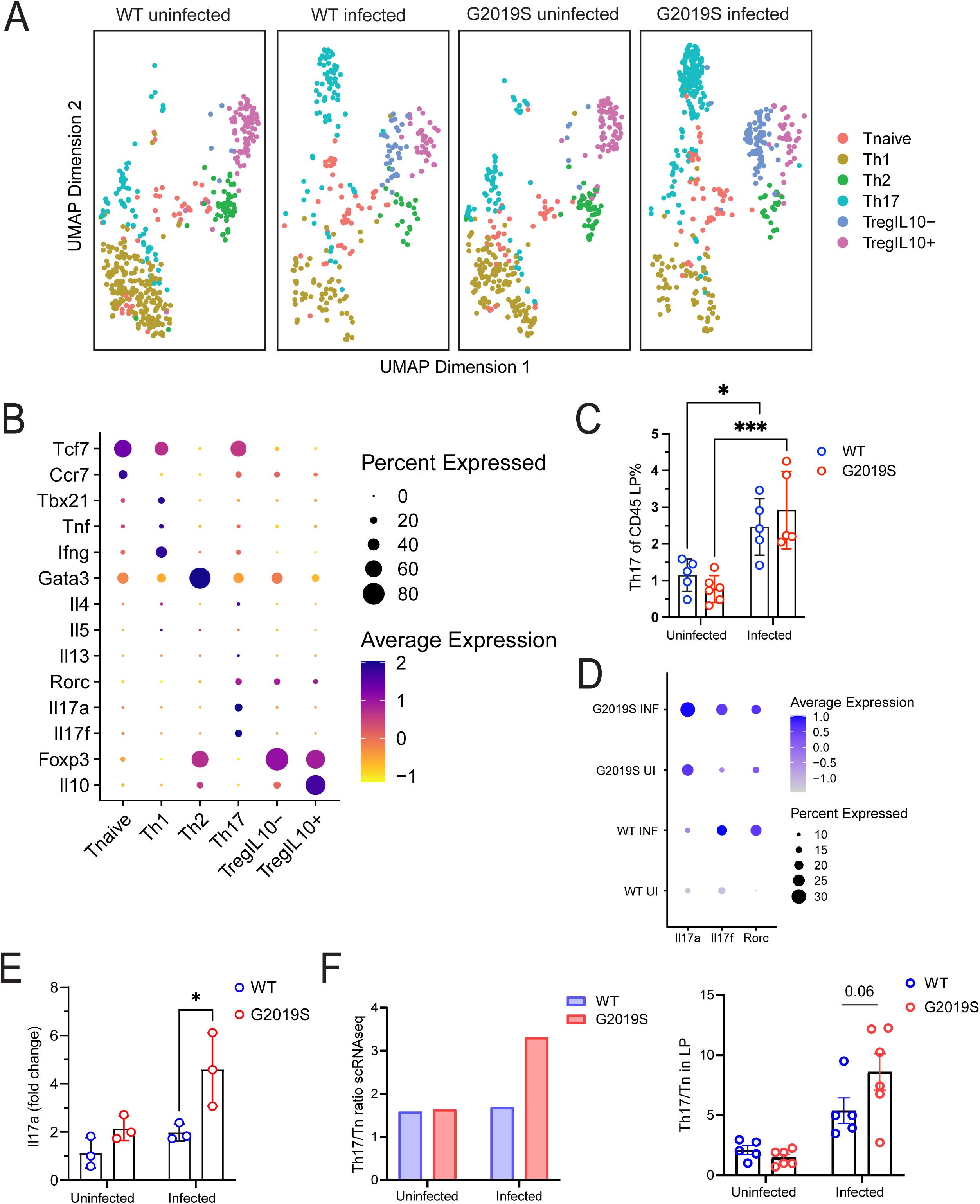
Colonic infection in LRRK2 G2019S mice initiates a skewing towards Th17 CD4^+^ T cells. (A) Re-clustering of CD4 T cell population based on effector T cell identities, depicted in a UMAP plot. (B) CD4 T cell subset markers used to identify effector cells based on literature. (C) Percent Th17 (RORgt+) CD4 T cells of CD45+ cells isolated from the colonic lamina propria across conditions. Data are presented as mean ± SD and analyzed by two-way ANOVA with Sidak post-test. * p< 0.05, ** p<0.01, p < 0.001, p<0.0001. n=5-6 mice per group. One independent experiment is presented. (D) Dotplot depicting average expression of *Il17a*, *Il17f,* and *Rorc* transcripts within Th17 effector subset. (E) *Il17a* expression fold change of sorted CD45+ immune cells from the lamina propria measured through quantitative PCR (qPCR). Data are presented as mean ± SD and analyzed by two-way ANOVA with Sidak post-test. * p< 0.05. n=3 mice per group. One independent experiment is presented. All qRT-PCR data was normalized to *Gapdh* using the ΔΔCt method. (F) Left: Ratio of absolute counts of Th17 effector cells over T naïve cells in scRNAseq dataset. Right: Ratio of Th17 effector cells over T naïve cells in CD45+ lamina propria populations in flow cytometry data.

## Discussion

LRRK2 is highly expressed in immune cells, and its mutations are common risk factors for both PD and IBD, placing LRRK2 at the intersection of the immune system, the intestine, and PD (de Guilhem de Lataillade et al., 2023; Herrick & Tansey, 2021; Kim et al., 2018; Lee et al., 2021; Liu et al., 2017). In the present study, we have systematically investigated the early immune response to intestinal infection in the context of the PD-associated mutation – LRRK2 G2019S. At day 7 post *C. rodentium* infection, we observed a marked accumulation of neutrophils in LRRK2 G2019S mouse colons compared to WT colons. This coincided with elevated tissue pathology and inflammation driven by increased erosion, edema, and goblet cell loss scores. Functionally, LRRK2 G2019S neutrophils showed cell-intrinsic increases in degranulation, chemotaxis, and NETosis under inflammatory conditions compared to WT controls. Moreover, infected LRRK2 G2019S mice demonstrated skewing of CD4 subsets towards a Th17 profile, with increased transcriptional levels of *Il-17a*. Together these results point to a role for *LRRK2* in immune cells during the earliest responses following intestinal infection.

With respect to the LRRK2 G2019S mutation, most studies to date have focused on its role in monocytes, macrophages and microglia (Kim et al., 2018; Liu et al., 2017; Moehle et al., 2015; Navarro et al., 2024; Nazish et al., 2023; Wallings et al., 2023). However, a few studies have investigated the effects of G2019S mutation in the context of neutrophils. Here, we demonstrate that under infection conditions the G2019S mutation not only causes increased presence of neutrophils in the lamina propria, but transcriptionally, these neutrophils have a greater pro-inflammatory type I and II IFN response compared to WT mice. This is consistent with previous studies, where the G2019S mutation has been associated with increased IFN signaling under inflammatory conditions (Yang et al., 2024). Moreover, LRRK2 expression has been consistently shown to be induced by IFN-γ stimulation (Gardet et al., 2010; Hakimi et al., 2011; Panagiotakopoulou et al., 2020; Thévenet et al., 2011), which is further supported by the presence of IFN response elements in the LRRK2 promoter region (Gardet et al., 2010). Taken together, there is a close link between LRRK2 and IFN pro-inflammatory pathways which is relevant in the context of PD.

Functionally, we also demonstrate that LRRK2 G2019S neutrophils *in vitro* have an increased ability to migrate, degranulate, and undergo NETosis, in a cell-intrinsic manner. They underwent more migration in response to the chemoattractant fMLP and *C. rodentium*, which was further supported *in vivo* by upregulated chemotaxis related DEGs in infected G2019S neutrophils (*Plaur, Cxcl10, Ccl6, Ccr1*). LRRK2 kinase activity and G2019S-mediated alterations in migratory function have previously been characterized in microglia, macrophages, and in a previous study with neutrophils with conflicting results (Mazaki et al., 2023; Moehle et al., 2015; Panagiotakopoulou et al., 2020). Reports by Moehle et al. and Panagiotakopoulou et al. indicate that G2019S mutation induces increased migration in myeloid cells and iPSC-derived microglia, respectively (Moehle et al., 2015; Panagiotakopoulou et al., 2020). Moehle et al. demonstrated that myeloid cells with G2019S mutations have enhanced chemotaxis towards inflammatory stimulus thioglycolate in both *in vitro* migration assays and *in vivo* migration of myeloid cells into the peritoneum following thioglycolate injection (Moehle et al., 2015). Similarly, Panagiotakopoulou et al. reported increased migration and phagocytosis in iPSC-derived microglia carrying the G2019S mutation, showing increased mobility of microglia towards adenosine triphosphate as a chemoattractant (Panagiotakopoulou et al., 2020). Results from Mazaki et al. in dH-L60-cell line derived neutrophils indicated LRRK2 knockdown decreased neutrophil migration towards fMLP, whereas inhibition of kinase activity with an inhibitor increased their migratory capacity (Mazaki et al., 2023). These three papers proposed varying mechanisms of LRRK2-mediated migration through binding of LRRK2 to actin-regulatory proteins or binding to MFN2 on the outer mitochondrial membrane. Notably, we uncovered a striking difference in actin cytoskeleton organization and cell shape in LRRK2 G2019S neutrophils exposed to *C. rodentium* (Fig. 5). This is consistent with previous reports of LRRK2 binding to actin regulatory proteins and our own scRNAseq results uncovering differential expression of actin cytoskeleton-related genes (*Actg1*) (Moehle et al., 2015). With respect to PD, several studies and meta-analyses have indicated the association between increased neutrophil to lymphocyte ratio (NLR) in people with PD compared to healthy controls (Grillo et al., 2023; Hosseini et al., 2023), providing further evidence of the critical role neutrophils may play in peripheral immunity and pathophysiology of PD. Future studies, aimed at discerning the exact mechanisms and consequences of these perturbations are currently underway. Overall, understanding the cellular function of LRRK2 within the neutrophils, and its implication both in PD and IBD can help us understand its pathophysiological role in these two diseases.

scRNAseq and flow cytometry data at day 7 post *C. rodentium* infection revealed a skewing of CD4^+^ T cells towards Th17 effectors in G2019S mice compared to WT, accompanied by significantly upregulated IL-17A expression in sorted CD45^+^ lamina propria cells. Neutrophils and Th17 cells communicate in a bidirectional manner, but the specific dynamics of their interaction mediated by G2019S during infection remains unclear. Th17 cells are potent recruiters of neutrophils, indirectly recruiting them to sites of inflammation via IL-17A. IL-17A induces the epithelial, endothelial, and stromal cells to produce IL-1, IL-6, TNF, CXCL8 (CXCL1 in mice), G-CSF, GM-CSF, which are neutrophil chemoattractants (Fan et al., 2023). However, neutrophils can regulate recruitment of Th17 cells through release of CCL20, CCL2, and CXCL10 in response to IFNγ and LPS (Pelletier et al., 2010). Additionally, neutrophil NETosis has been reported to cause skewing of T naïve cells into Th17 cells in a dose dependent manner (Wilson et al., 2022). Histones that decorate released NETs can activate toll-like receptor 2 in naïve T cells, which will phosphorylate STAT3 and facilitate Th17 differentiation (Wilson et al., 2022). According to *C. rodentium* infection kinetics, neutrophils are one of the first immune cells recruited to the colonic lamina propria, followed by T cells, including Th17 cells (Mullineaux-Sanders et al., 2019). Therefore, we hypothesize that in G2019S mice, neutrophil presence is initially increased, followed by NETosis, which could promote increased Th17 differentiation and play into a positive feedback loop with neutrophil recruitment.

However, to clarify the sequence of events, further analyses are needed. Park et al. previously showed that LRRK2 G2019S in rats caused increased presence of immature myeloid cells with suppressive activity on Th17 cell differentiation in the gut in response to acute (trinitrobenzene sulfonic acid (TNBS)-induced) and chronic (DSS-induced) colitis (Park et al., 2017).However, in comparison to our work, *C. rodentium* infection itself induces a much more robust Th17 response compared to DSS-induced colitis, indicating that LRRK2 G2019S mutation in the context of Th17 skewing may be dependent on the inflammatory setting.

Understanding the interplay of gene-environment interactions is critical to uncover the multifaceted etiology of PD. We previously showed that repeated intestinal infection with *C. rodentium* in mice harbouring a mutation in another PD-related gene (*Pink1^-/-^)* triggered the development of PD-relevant dopamine-sensitive motor symptoms later in life. The study also revealed the infection-initiated presentation of mitochondrial antigens by antigen presenting cells, which triggered the development of cytotoxic CD8+ T lymphocytes that are specific to mitochondria in both the brain and the periphery (Matheoud et al., 2019). Recently, a study from our research group mapped the early immunological events at the site of *C. rodentium* infection in *Pink1* ^-/-^ mice. They uncovered that at the peak of the innate immune response to *C. rodentium* (1-week post infection), *Pink1*^-/-^ mice had dysregulated differentially expressed genes within myeloid cells of the monocyte/macrophage lineage (Recinto et al., 2024). Specifically, during infection, *Pink1*^-/-^ monocytes skewed towards “mature” macrophage- and DC-like profiles, with an increased capacity of MHC II presentation (Recinto et al., 2024). Like *Pink1* ^-/-^, we also demonstrate that the monocytes in LRRK2 G2019S mice were affected 1-week post *C. rodentium* infection. However, our results in LRRK2 G2019S mice showed increased immature- Ly6C^High^ monocytes in the colonic lamina propria, with downregulation of MHC II encoding genes and MHC II surface protein levels (data not shown), distinct from the differences seen in the *Pink1^-/-^* mice. Although the mechanisms regulating monocytes differ between LRRK2 G2019S (a common genetic variant linked to an increased risk of PD) and loss-of-function mutations in PINK1 (rare mutations associated with familial and early-onset PD) (Day & Mullin, 2021), our collective work highlights the involvement of two distinct PD-related genes in reprogramming of the immune response to infection.

It is becoming increasingly apparent that chronic intestinal inflammation could play a key role in early, prodromal pathophysiology of PD. Notably, drawing similarities between IBD and PD could help us elucidate underlying mechanisms, as they are both chronic progressive diseases with complex etiology involving genetic and environmental factors (Lee et al., 2021). IBD is reported to increase risk for PD development by approximately 20% (Lee et al., 2021; Li et al., 2023; Park et al., 2019; Villumsen et al., 2019). Notably, people with IBD receiving anti-TNFα treatment exhibit a reduced risk of developing PD compared to those not on anti-TNFα (Peter et al., 2018). Involvement of pathogenic LRRK2 mutations in the kinase domain is relevant to both diseases, whereby G2019S causes increased risk for PD and N2081D is associated with Crohn’s Disease (CD – one form of IBD) and PD (Herrick & Tansey, 2021). Both mutations lead to increased kinase activity and phosphorylation of the LRRK2 substrate Rab10 (Hui et al., 2018; Thévenet et al., 2011). Expression of LRRK2 itself is also elevated in biopsies from CD patients (Gardet et al., 2010). Additionally, relevant to our findings, IBD studies have shown that dysregulated neutrophilic activation and accumulation can also exacerbate disease, causing chronic inflammation and mucosal injury (Cassatella et al., 2019; Friedrich et al., 2021; Pelletier et al., 2010; Sanchez-Garrido et al., 2024).

While we are characterizing the earliest colonic responses to infectious colitis in the G2019S mouse model, others have explored how LRRK2 G2019S impacts DSS-induced intestinal inflammation (Cabezudo et al., 2023; Fang et al., 2024; Lin et al., 2022). Cabezudo et al. demonstrate that G2019S mutants under DSS treatment have significantly elevated colonic histopathological score compared to WT DSS, coupled with intense infiltration of leukocyte populations, and of note, increased presence of MPO+ cells. Using WT bone marrow transplantation into G2019S mice, they could fully rescue the inflammatory phenotype. Likewise, inhibition of LRRK2 G2019S kinase activation could partially rescue the phenotype. This indicates the clear involvement of LRRK2 kinase activity in immune cells in exacerbation of inflammatory phenotypes in G2019S mice during DSS-induced colitis. Similarly, Fang et al. and Lin et al. also demonstrated that G2019S mice were more vulnerable to DSS induced colitis, both with subsequent inflammatory consequences (Fang et al., 2024; Lin et al., 2022). While Fang et al. showed increased macrophage frequency to the colonic lamina propria, as well as increased alpha-synuclein presence, Lin et al. showed increased intestinal expression of PRRs like TLRs, NF-kβ, and pro-inflammatory cytokine TNF-α. Our work extends these findings through global assessment of the early response to a self-limiting infection with a natural mouse intestinal pathogen. This work reveals that LRRK2 G2019S impacts the very early response to infection in the gut, and that amongst all cells of the lamina propria, neutrophils are arguably the most prominently affected by the mutation in this setting. Furthermore, we uncover a number of cell-intrinsic differences in LRRK2 G2019S neutrophils, suggesting that these cells may play a primary role in the overall immune dysfunction observed in the infected LRRK2 mutant mice.

Others have demonstrated that chronic, prolonged treatment of LRRK2 G2019S mice with DSS lead to development of PD-like motor impairments (Fang et al., 2024; Lin et al., 2022) and that DSS coupled with injection of a human alpha-synuclein overexpression vector (recombinant adeno-associated virus) led to PD-like neurodegeneration in G2019S mice (Cabezudo et al., 2023). Notably, we did not observe any development of motor symptoms in the LRRK2 G2019S or WT mice within the time frame of our experiments. This is not surprising since both in our previous studies in *Pink^-/-^*mice and in the LRRK2 G2019S DSS colitis studies mentioned above, motor symptom development required months of time and multiple exposures to infectious or inflammatory stimuli. While beyond the scope of this paper, which focuses on the impact of LRRK2 very early on in the prodromal phase, it would be of interest to determine if repeated exposures to *C. rodentium* and/or aging of the mice would lead to development of PD-like motor symptoms.

Taken together, our results demonstrated that the LRRK2 G2019S mutation has a profound impact on the early response to gut infection. This is evident by the increase of neutrophil migration, dysfunctional degranulation, increase in NETosis, inflammatory-mediated tissue damage, and elevation of IL-17A expression. Considering that immune dysregulation plays a role in the development and progression of neurodegenerative disease, our findings could contribute to a better understanding of mechanisms within the prodromal phase of PD. This understanding could inform the development of pharmacological targets for early detection and intervention in PD.

## Supplemental figure legends

Supplemental Figure 1. LRRK2 G2019S mice have similar control of C. rodentium infection with minor changes in fecal water composition. Male and female LRRK2 G2019S and WT mice were infected once with approximately 1x10^9^ CFU of *C. rodentium*. (A) From days 0-7 of infection, weight change was measured. Data are represented as individual mouse points with the average weight loss trend per group. Depicted is one representative of two experiments. n=5-6 mice per group. On day 5 or 7 of infection, mice were placed individually in cages without bedding, food, or water for quantification of the number of stools excreted by each animal and their water content. (B) Number of fecal pellets per hour. Data are represented as mean ± SD. Depicted is one representative graph of two experiments. n=6-9 mice per group. (C) Water content of feces. Data are represented as mean ± SD and analyzed by two-way ANOVA with Sidak post-test. *p<0.05. Two pooled experiments, n=12-13 mice per group.

Supplemental Figure 2. LRRK2 G2019S infected mice have increased presence of Ly6c^high^ monocytes with dysregulated differentially expressed gene profiles. (A) Pathway enrichment analysis completed on DEGs obtained by comparing G2019S and WT infected monocytes. DEGs were run against Gene ontology (GO) biological pathways database using *ClusterProfiler.* P value cut off = 0.05, q value cut off = 0.2. Top 30 GO term categories depicted as a Treeplot (Left). Cnetplots depict association of differentially expressed gene hits, and which GO term pathways they correspond to (Right). (B) Percentage of Ly6c^High^ monocytes of lamina propria assessed through flow cytometry. Data are presented as mean ± SD and analyzed by two-way ANOVA with Sidak post-test. * p< 0.05. n=5-6 mice per group. One independent experiment is presented. (C) Subclustering of monocytes and annotated as Ly6c2^high^, Cx3cr1^+^ (differentiating towards Macrophage-like), or Itgax^+^ (differentiating towards DC-like). (D) Proportional analysis conducted through permutation test comparing G2019S infected and WT infected conditions. Significance cut-off: FDR < 0.05.

Supplemental Figure 3. LRRK2 G2019S infected mice have dysregulated differentially expressed gene profiles in γδ T Cells. (A) Pathway enrichment analysis completed on DEGs obtained by comparing G2019S and WT infected γδ T Cells. DEGs were run against Gene ontology (GO) biological pathways database using *ClusterProfiler.* P value cut off = 0.05, q value cut off = 0.2. Top 30 GO term categories depicted as a Treeplot (Left). Cnetplots depict association of differentially expressed gene hits, and which GO term pathways they correspond to (Right).

Supplemental Figure 4. Colon histopathology score is affected by LRRK2 G2019S mutation after *C. rodentium* infection. Male and female LRRK2 G2019S and WT mice were infected once with approximately 1x10^9^ CFU of *C. rodentium*. Colons were harvested 7 days after infection, fixed in 10% formalin for 24 h, washed in 70% ethanol, and diaphanized in xylene before embedding in paraffin. Sections were stained with Hematoxylin and Eosin (H&E) and scored for histopathology. Histopathological scoring was conducted blindly by an expert veterinary pathologist based on the scoring criteria of colon lesions: inflammatory infiltrate (0- 4), polymorphonuclear (PMN) cell infiltrate (0-4), loss of crypts (0-2), proportional loss of goblet cells (0-2), edema (0-1), erosion or ulceration (0-3), hemorrhage (0-2), and necrosis (0-1). Total pathology score (0-19) was the sum of all individual lesion scores. Data are presented as mean ± SD and analyzed by two-way ANOVA with Sidak post-test. * p< 0.05 and *** p <0.001. n= 7-11 mice per group. Two independent experiments are presented as a pool.

## Author contributions

JP completed scRNAseq bioinformatic analysis with guidance from SR. NM led *in vivo* experiments. NM performed flow cytometric staining and analysis of the colonic lamina propria cells with assistance from AK. JP, NM, SR, and AK participated in processing cells for scRNAseq. CMQ completed histopathological scoring. ZL, KC, and AT troubleshot and executed *in vitro* migration and NETosis assays. NM and AK completed degranulation and MPO assay. CR bred and maintained animal colony under the supervision of AJM. SW contributed to analysis of neutrophils *in vivo* under the supervision of ILK. JP and NM prepared figures, drafted, and edited the manuscript. MD provided feedback on the project. SG and JAS supervised the project, edited the manuscript, and approved the final draft. All authors contributed and approved the submitted version.

