## Supplemental Figure 1 for "LRRK2 G2019S mutation incites increased cell-intrinsic neutrophil effector functions and intestinal inflammation in a model of infectious colitis"

Supplemental Figure 1. LRRK2 G2019S mice have similar control of *C. rodentium* infection with minor changes in fecal composition and frequency of evacuation.

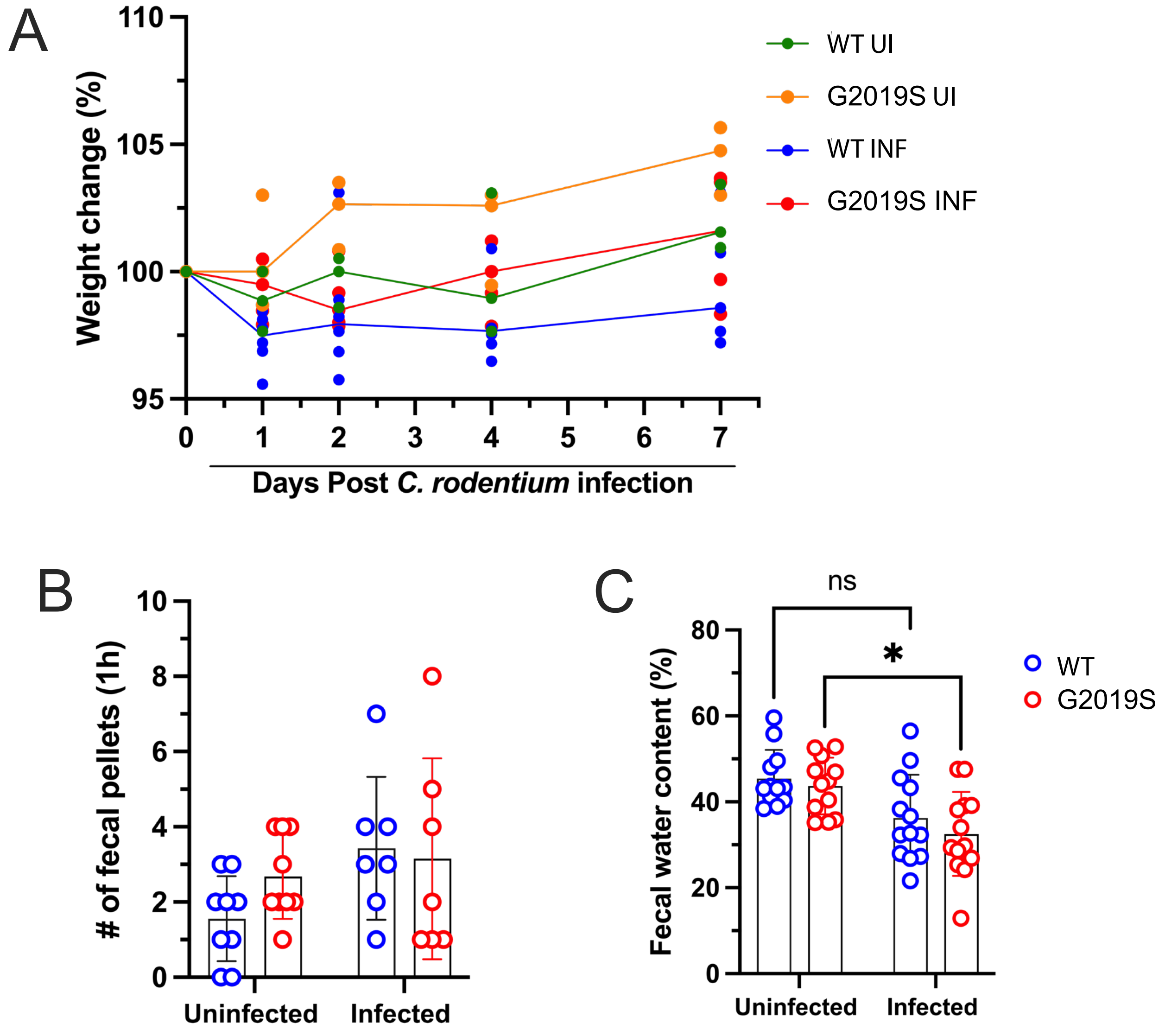
