## Supplemental Figure 2 for "LRRK2 G2019S mutation incites increased cell-intrinsic neutrophil effector functions and intestinal inflammation in a model of infectious colitis"

Supplemental Figure 2. LRRK2 G2019S infected mice have increased presence of Ly6c$^{high}$ monocytes with dysregulated differentially expressed gene profiles.

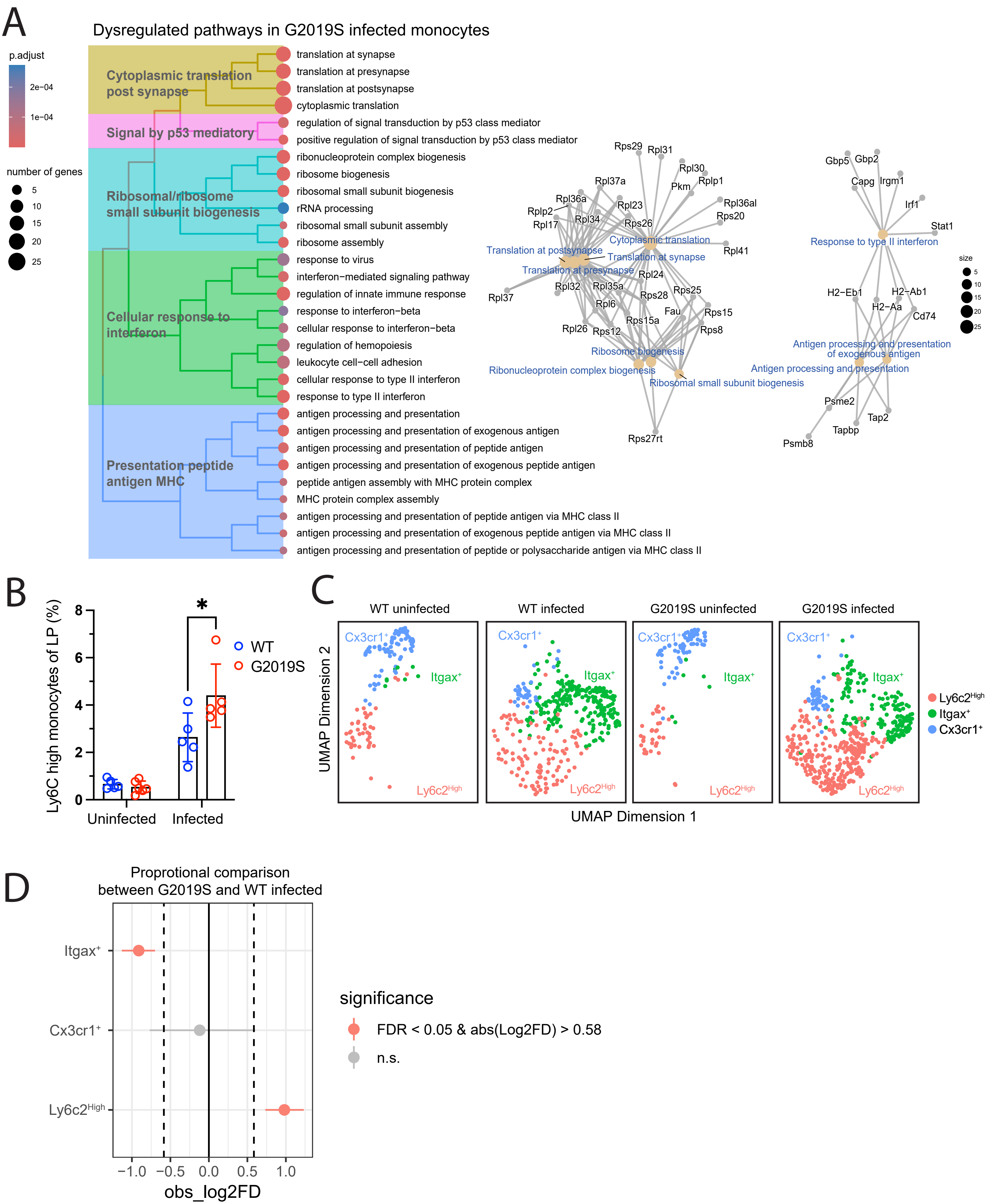
