## Supplemental Figure 3 for "LRRK2 G2019S mutation incites increased cell-intrinsic neutrophil effector functions and intestinal inflammation in a model of infectious colitis"

### Supplemental Figure 3. LRRK2 G2019S infected mice have dysregulated differentially expressed gene profiles in γδ T Cells.

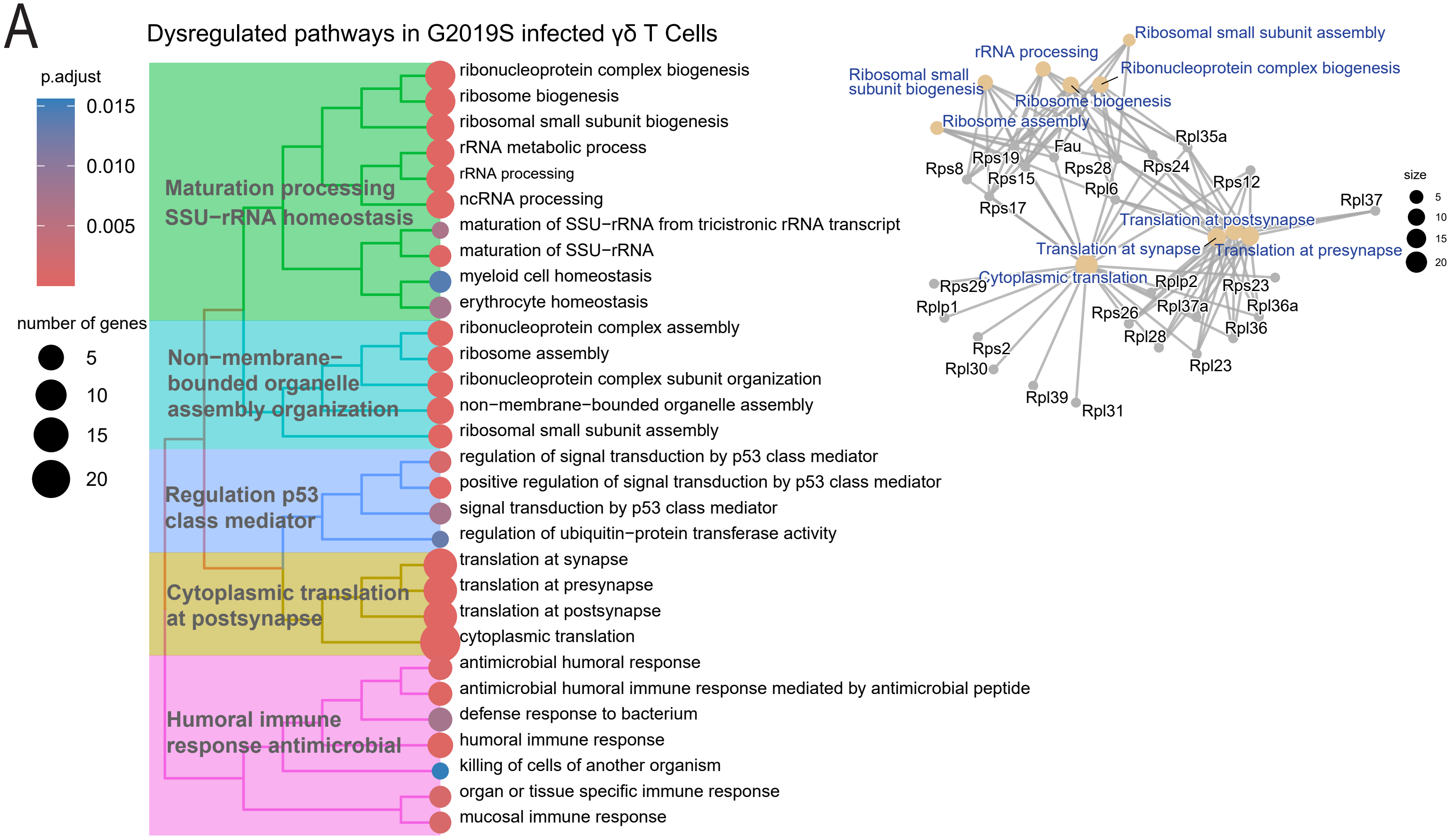
