## Supplemental Figure 4 for "LRRK2 G2019S mutation incites increased cell-intrinsic neutrophil effector functions and intestinal inflammation in a model of infectious colitis"

Supplemental Figure 4. Colon histopathology score is affected by LRRK2 G2019S mutation after *C. rodentium* infection.

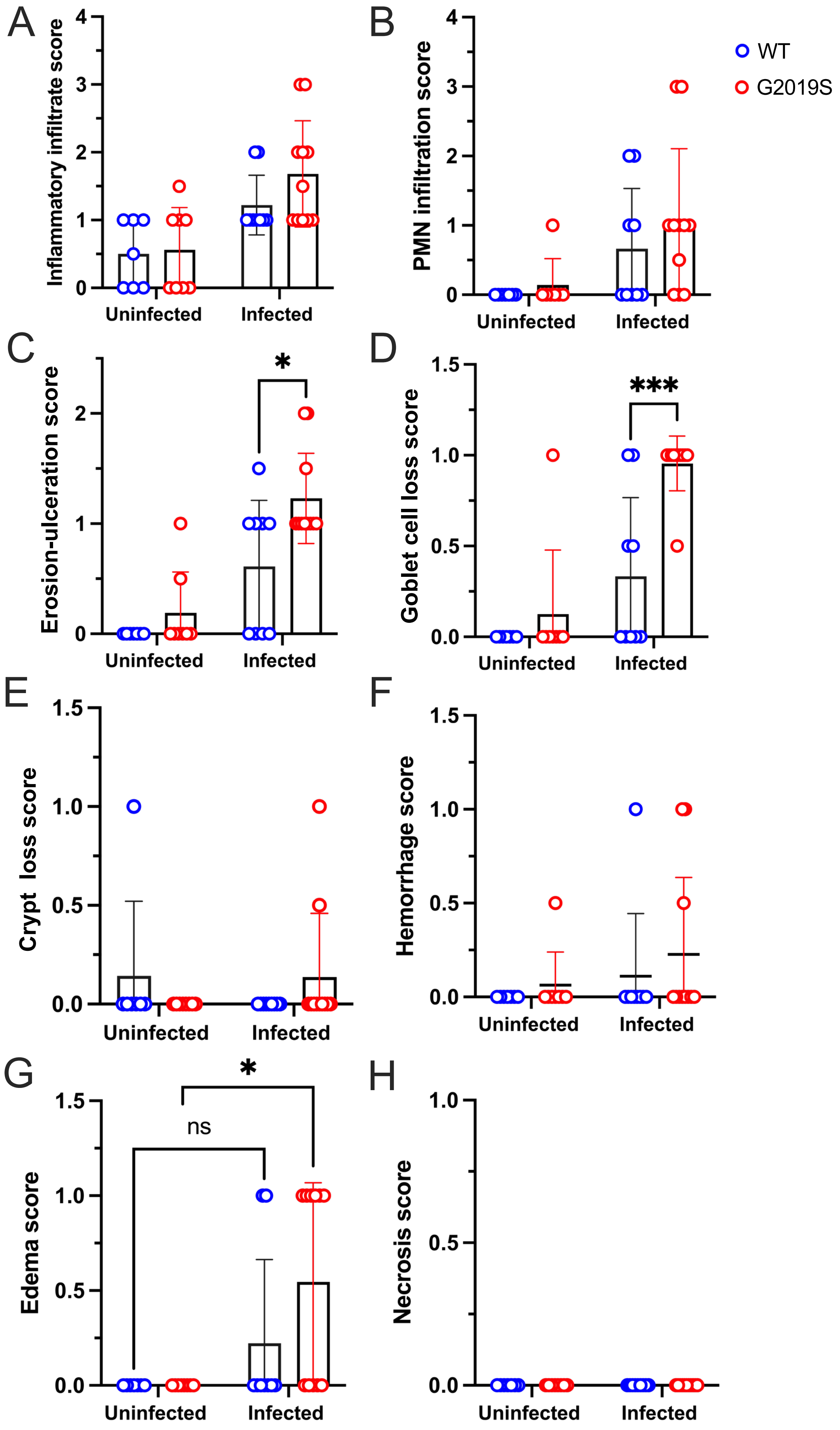
